# Nutritional history affects stress-resistance architecture across ageing and sex

**DOI:** 10.64898/2026.09.08.750110

**Authors:** Devashish Kumar, Mohankumar Chandrakanth, S Chetan, Chand Sura, Nishant Kumar, Sudipta Tung

## Abstract

Diet can influence lifespan and age-dependent physiological function, but how nutrition during development and adulthood affects relationships among stress resistance traits across age remains unclear. Using an isocaloric full-factorial larval × adult diet design in outbred *Drosophila melanogaster*, we tested how protein– and carbohydrate-biased diets affected starvation, desiccation, cold, osmotic, and heat resistance across the adult life course and sex. We also measured whole-body triacylglyceride, protein, body weight, and water content, and compared responses with matched lifespan and reproductive-output data. Resistance declined across the adult life course, but the traits did not vary as a single phenotype: among the traits included in the covariance analysis, desiccation, osmotic, and heat resistance formed a tightly correlated core, whereas starvation resistance remained comparatively peripheral in both sexes. Developmental diet produced trait-specific effects, whereas protein-biased adult diet favoured heat and osmotic resistance, paralleling effects on lifespan and reproductive output, but had opposite effects on starvation and cold resistance in specific contexts. Thus, nutrition alters rather than uniformly improves stress resistance, and covariance among resistance traits does not necessarily predict how those traits respond to nutritional perturbation.

## Introduction

Ageing is accompanied by progressive loss of somatic maintenance and functional performance, reducing the capacity to withstand environmental challenge and impacting survival and reproduction in late life ^1,2^. Assays of performance under environmental stress, including nutritional, osmotic, thermal, and desiccation challenges, provide tractable functional readouts of stress tolerance and, when measured across age, offer a multivariate view of age-dependent stress performance ^3,4^. Stress resistance is also frequently associated with longevity in experimental systems, although the relationship is trait– and context-dependent ^5–7^. Such resistance is unlikely to be cost-free: stress defence draws on energetic and material reserves, requires their mobilization under challenge, and must be balanced against allocation to other functions such as somatic maintenance and reproduction ^3,8^. Because nutrition supplies the material and energetic basis for these processes, understanding how diet organizes stress resistance across age is central to explaining how nutritional interventions shape physiological function and life-history traits.

Nutritional geometry has provided a powerful framework for mapping how macronutrient balance affects lifespan and reproduction. In *Drosophila*, mice, and other systems, protein and carbohydrate balance can generate distinct nutrient optima for longevity and fecundity, with reproduction often favoured by higher protein availability and lifespan favoured by lower-protein, carbohydrate-biased diets that do not lead to malnutrition ^9–11^. This relationship is context-dependent rather than fixed: targeted amino-acid manipulation can separate effects on fecundity and lifespan ^12^, and recent work in an outbred *Drosophila melanogaster* population showed that protein-biased diets can increase both fecundity and lifespan under isocaloric macronutrient manipulation ^13^. In parallel, stress-resistance studies have asked whether tolerance to different stressors reflects a generalized stress-resistance syndrome or stress-specific physiology, with evidence for both correlated responses among stressors and trait-specific or sex-specific trade-offs ^5,14,15^. These two research themes have rarely been integrated: nutritional studies of ageing typically emphasize lifespan and reproduction, whereas stress-resistance studies less often map broad resistance suites under controlled macronutrient manipulation across life stage, age, and sex. This leaves unresolved whether stress-resistance traits that covary across conditions are also those that respond similarly to nutritional perturbation.

Stress-resistance traits that covary across conditions may be interpreted as phenotypically integrated, because covariance among traits is widely used to characterize integration among potentially functionally related traits ^16,17^. However, covariance and response to perturbation are analytically distinct ^18,19^: a correlation across age, sex, and diet conditions does not imply that a specific nutritional contrast will change those traits in the same direction. Conversely, traits weakly coupled in the overall covariance structure may respond in parallel when diet is manipulated. We therefore distinguish two complementary levels of phenotypic organization, using “architecture” here to describe relationships among measured traits rather than their underlying genetic basis: a covariance architecture, describing which traits vary together across conditions, and a dietary-effect architecture, describing which traits show shared or opposing responses to nutrition. This distinction matters for ageing biology because trait correlations need not predict how the same traits respond to dietary perturbation or how those responses align with longevity and reproduction.

This problem of reconciling covariance and dietary-effect structure becomes especially complex when nutrition varies across the life course. In holometabolous insects such as *Drosophila*, larval and adult diets act on different physiological problems: larval nutrition shapes growth rate, developmental timing, body size, and organ growth through nutrient-sensitive endocrine pathways ^20,21^, whereas adult nutrition supplies the immediate resource environment for somatic maintenance, reproduction, and stress defence ^9,22,23^. Developmental nutrition can carry over into adult function, influencing ovarian maturation, early reproduction, and starvation resistance, indicating that adult stress performance is partly shaped by larval nutritional history ^24–27^. Resolving how developmental and adult diet each contribute to age-dependent resistance, additively, stage-specifically, or through their interaction, requires manipulating the two diets independently rather than assuming that developmental and adult nutrition act concordantly.

Here, we used an isocaloric full-factorial larval × adult diet design in a large outbred population of *Drosophila melanogaster* to test how protein-biased and carbohydrate-biased nutrition during development and adulthood partially organize adult stress resistance across age and sex. The diets were formulated to equal caloric density but differed in P:C ratio, and were independently manipulated during development and adulthood, allowing larval, adult, and larval × adult effects to be separated. We assayed five stress-resistance traits — starvation resistance, desiccation resistance, cold resistance, osmotic stress resistance, and heat resistance — at three post-egg-collection sampling points spanning early to later adult life in both sexes. We first examined the covariance architecture of age-dependent stress resistance, then tested how developmental and adult diets modified individual traits, and finally asked whether patterns of diet-induced responses across resistance traits corresponded to their covariance structure and how these responses related to matched longevity, fertility, and physiological measurements.

## Results

### Stress resistance forms an age-structured covariance architecture shared across sexes

To define the covariance architecture of stress resistance, we examined the four resistance traits summarized on directly comparable time-to-event scales: starvation resistance (SR), desiccation resistance (DR), osmotic/high-salt stress resistance (OSR), and heat resistance (HR). Condition-level values were standardized before PCA and correlation analyses (see Methods). This allowed us to compare covariation among traits measured in different units across the full larval × adult diet design. Cold resistance (CR), which was measured using sex– and age-specific calibrated challenge durations, was retained for trait-specific and dietary-response analyses but excluded from cross-stratum covariance analyses.

Sex-specific PCA showed a highly similar multivariate structure in females and males. PC1 captured 87.3% of the variance in females and 87.5% in males, while PC2 explained 9.9% and 9.5%, respectively (Figure 2A,B). In both sexes, conditions were strongly ordered along PC1 across the adult life course, with early-life conditions clearly separated from mid– and later-life conditions across dietary regimes, while mid– and later-life conditions were closer and partly overlapping (Figure 2A,B). This pattern was supported by analysis of PC1 scores, which differed significantly among life-course points after accounting for dietary regime (females: *F*_2,6_ = 580.13, *P* = 1.36 × 10^−7^; males: *F*_2,6_ = 117.42, *P* = 1.55 × 10^−5^). No corresponding age effects were detected for PC2 (females: *P* = 0.284; males: *P* = 0.809). Thus, progression across the adult life course primarily structured the dominant multivariate resistance axis, with the visual separation suggesting that the largest shift occurred between early and mid-life.

**Figure 1.**
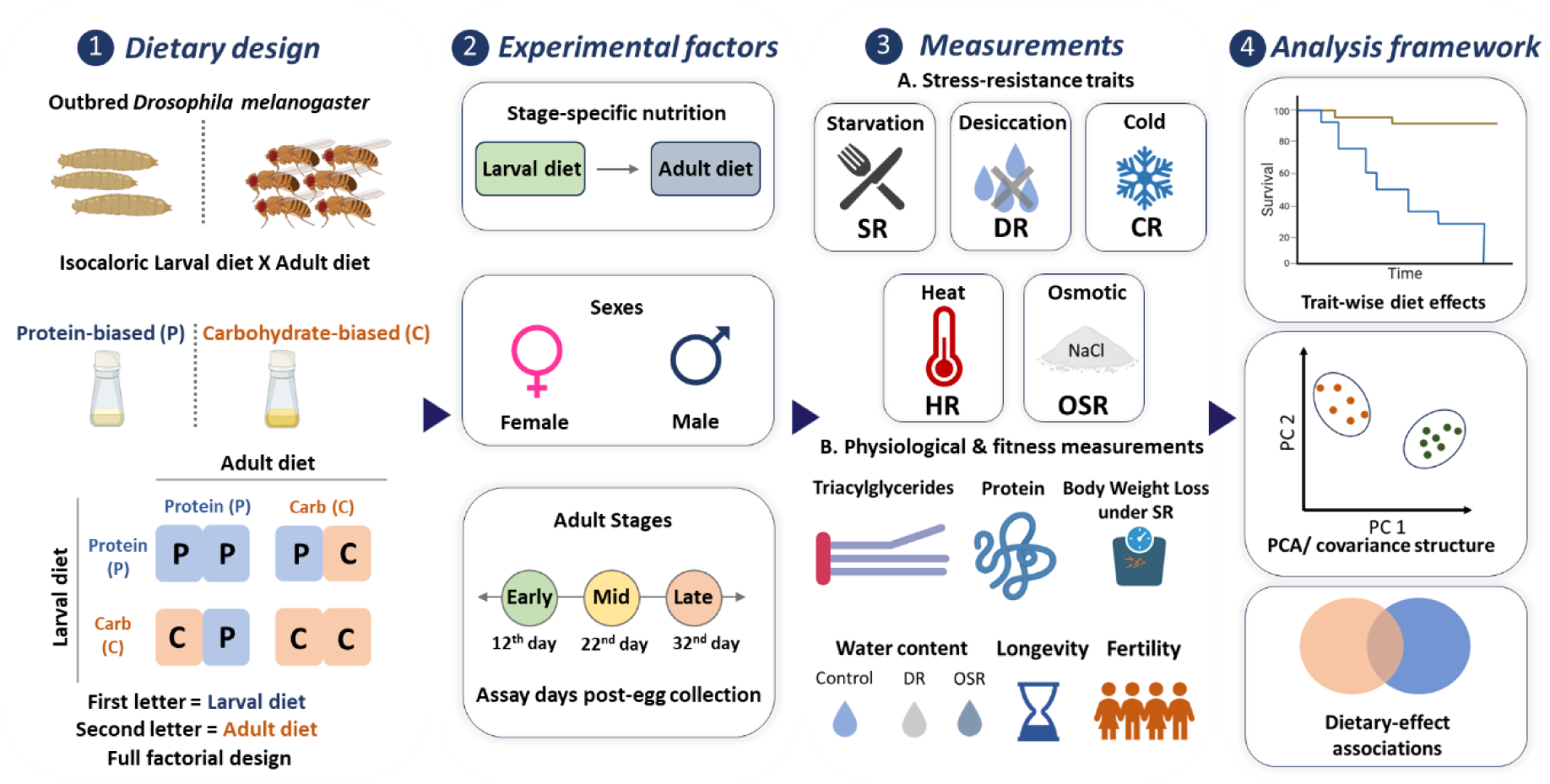
Experimental design and analytical framework. Overview of the isocaloric full-factorial larval × adult diet design used to test stage-specific nutritional effects on stress resistance in an outbred *Drosophila melanogaster* population. Protein-biased (P) and carbohydrate-biased (C) diets were independently manipulated during larval and adult stages to generate PP, PC, CP and CC regimes, where the first letter denotes larval diet and the second denotes adult diet. Starvation resistance (SR), desiccation resistance (DR), cold resistance (CR), heat resistance (HR) and osmotic stress resistance (OSR) were assayed across sex and adult life course, alongside physiological and fitness-associated measurements. These data were integrated to assess trait-wise diet effects, covariance structure and dietary-effect associations with longevity and fertility.

**Figure 2.**
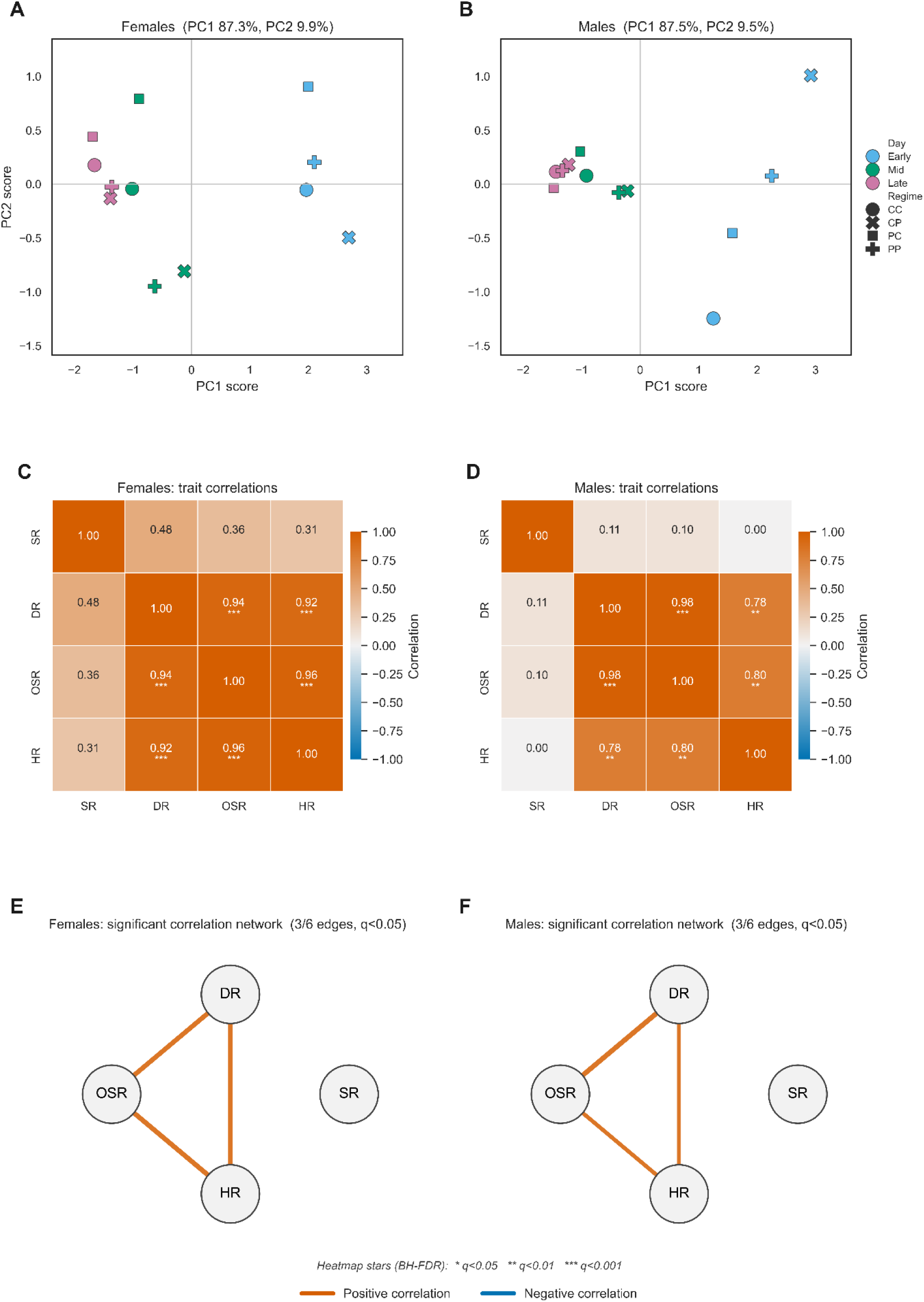
Stress resistance shows an age-structured covariance architecture shared across sexes. **A,B**, Principal component analysis of standardized condition-level SR, DR, OSR, and HR values in females (A) and males (B). Each point represents one life-course × dietary-regime condition. Colours indicate life-course point (early, mid and late) and symbols indicate larval × adult dietary regime (CC, CP, PC and PP). C,D, Sex-specific Pearson correlation matrices among SR, DR, OSR, and HR across condition-level means. Values indicate correlation coefficients; asterisks denote BH–FDR-adjusted significance levels: *q* < 0.05, ** *q* < 0.01, *** *q* < 0.001. E,F, FDR-filtered correlation networks showing significant correlations at *q* < 0.05. DR, OSR, and HR form a fully connected covariance core in both sexes, whereas SR remains peripheral.

Pairwise correlations showed a strongly structured covariance pattern (Figure 2C,D). All correlations were non-negative in both sexes, indicating no antagonistic covariance among the five traits. DR, OSR, and HR formed a tightly correlated core: DR–OSR correlations were near unity (females: *r* = 0.94; males: *r* = 0.98; both BH–FDR-adjusted *P* < 0.001), and both traits were strongly correlated with HR (females: DR–HR, *r* = 0.92; OSR–HR, *r* = 0.96; both adjusted *P* < 0.001; males: DR–HR, *r* = 0.78; OSR–HR, *r* = 0.80; both adjusted *P* < 0.01).

SR remained peripheral to this covariance core in both sexes. Its correlations with DR, OSR, and HR were positive but non-significant after FDR correction in females (*r* = 0.31–0.48) and weak in males (*r* = 0.00–0.11). Accordingly, FDR-filtered networks contained a fully connected DR–OSR–HR core in both sexes, whereas SR remained disconnected (Figure 2E,F).

A permutation-based integration analysis further showed that coupling among the four traits exceeded random expectation in both sexes (females: I = 0.39; males: I = 0.23; permutation *P* < 0.0001 and 0.0001 respectively; Supplementary Text S1 and Fig. S1).

Together, these analyses identify a covariance architecture largely shared across sexes: DR, OSR, and HR form a tightly correlated core, whereas SR remains comparatively peripheral. Because covariance describes which traits vary together but not how nutritional perturbation shifts them, we next examined dietary effects at the trait level.

### Developmental and adult diets modify stress resistance in trait-specific, stage-dependent ways

Having defined the covariance architecture, we next asked how developmental and adult nutrition modify the individual traits within it. For each trait, we used sex– and age-stratified mixed-effects models to evaluate the effects of larval diet, adult diet, and their interaction. We interpreted larval or adult main effects only where the larval × adult interaction was non-significant, and treated significant interactions as evidence that dietary effects depended on the specific larval–adult combination (Figure 3A; full sex– and age-stratified trait responses in Fig. S2-S6; statistical details in supplementary tables ST2-ST6).

**Figure 3.**
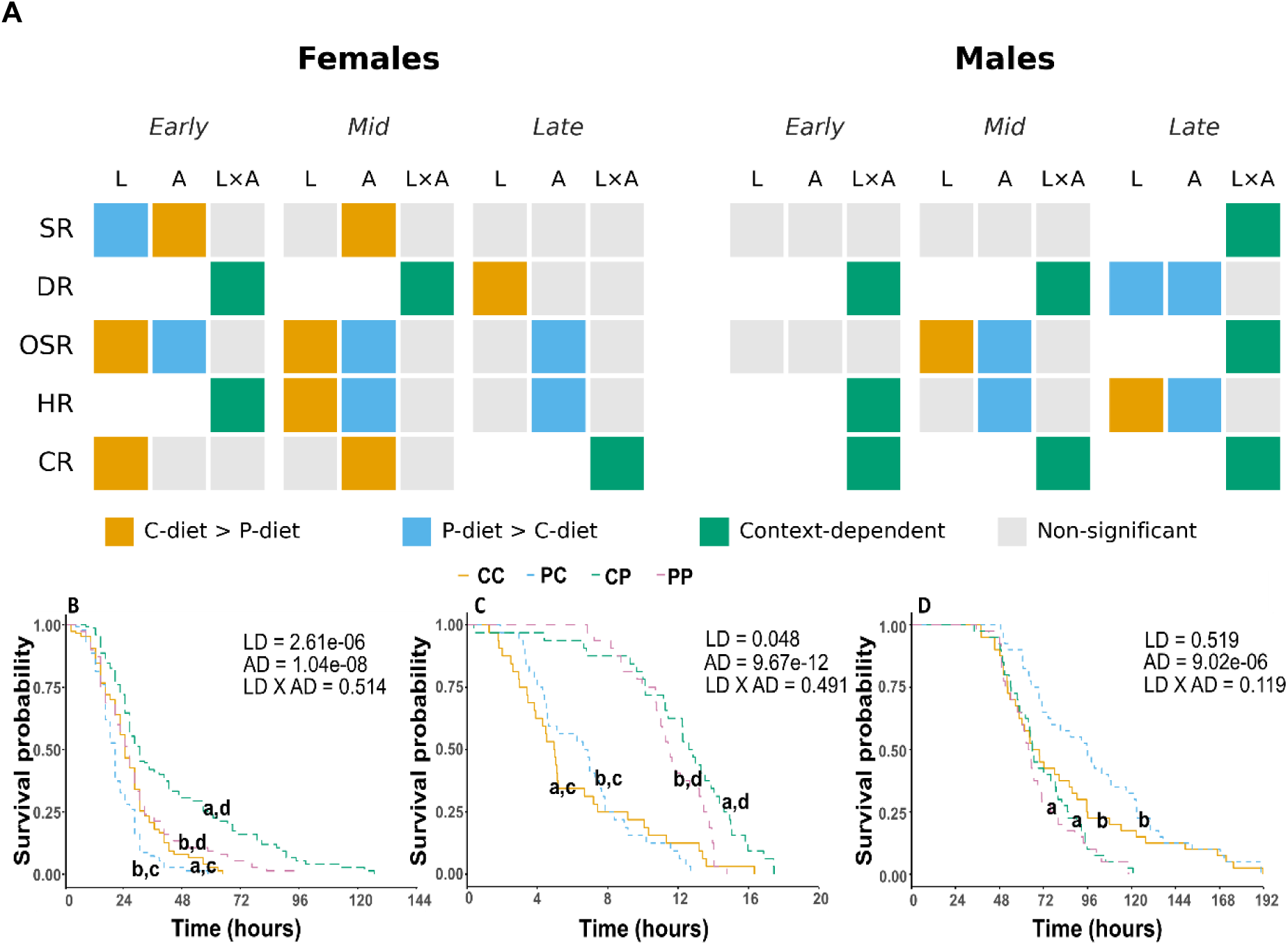
Stage-specific nutrition modifies stress resistance in a trait-, age– and sex-dependent manner. **A**, Summary of larval diet, adult diet and larval × adult diet effects on five stress-resistance traits across sex and age. Rows indicate starvation resistance (SR), desiccation resistance (DR), osmotic stress resistance (OSR), heat resistance (HR) and cold resistance (CR). Columns are grouped by sex and age class, with larval diet (L), adult diet (A) and larval × adult diet interaction (L × A) shown for each age. Colours indicate the direction or nature of significant dietary effects: gold, carbohydrate-biased diet > protein-biased diet; blue, protein-biased diet > carbohydrate-biased diet; green, significant context-dependent larval × adult diet effect; grey, non-significant effect. **B–D,** Representative survival curves illustrating trait-specific dietary responses in mid-age females; corresponding curves for all the sex × age strata are provided in Supplementary Figure S2-4. **B,** Osmotic stress resistance, showing significant larval and adult diet effects. **C,** Heat resistance, showing a weaker larval diet effect and a strong adult diet effect. **D,** Starvation resistance, showing a significant adult diet effect. Diet regimes are denoted by two letters, where the first letter indicates larval diet and the second indicates adult diet: CC, carbohydrate-biased larval and adult diet; PC, protein-biased larval and carbohydrate-biased adult diet; CP, carbohydrate-biased larval and protein-biased adult diet; PP, protein-biased larval and adult diet. LD, larval diet; AD, adult diet. P values indicate model-based tests of larval diet, adult diet and their interaction within the corresponding sex–age stratum. Different letters indicate statistical differences among diet regimes.

Adult diet produced the clearest recurrent directional effects, and these effects were trait-selective rather than uniform. Protein-biased adult diet increased HR across multiple sex–age strata (Mid F: P= 9.67 × 10⁻¹², Supp. Table-ST4c; Late F: P = 1.66 × 10⁻¹⁰, Supp. Table-ST4d; Mid M: P = 0.005, Supp. Table-ST4g; Late M: P = 7.01 × 10⁻⁴, Supp. Table-ST4h) and increased OSR wherever adult main effects were independently interpretable (Early F: P = 0.016, Supp. Table-ST3a; Mid F: P = 2.38 × 10⁻¹⁴, Supp. Table-ST3b; Late F: P = 0.005, Supp. Table-ST3c; Mid M: P < 2 × 10⁻¹⁶, Supp. Table-ST3e; Figure 3B, C). By contrast, carbohydrate-biased adult diet increased SR in early– and mid-age females (Early F: P = 0.026; Supp. Table-ST2a, Mid F: P = 9.0 × 10⁻⁶; Supp. Table-ST2b, Figure 3D). Thus, the same adult-diet contrast moved different resistance traits in opposite directions: adult-P favoured HR and OSR, whereas adult-C favoured SR in the female strata where significant main effects were detected. To assess whether the OSR pattern depended on differences in the basal assay-food matrix, we repeated the assay at day 22 post egg collection using a common P:C = 0.4 food supplemented with 4% NaCl across all dietary treatments. The treatment pattern was consistent with that obtained using the corresponding adult diets (Supplementary Fig. S12), indicating that the day-22 OSR differences were not driven solely by the assay-food matrix.

Larval diet produced fewer independently interpretable effects than adult diet, but the effects that were detected showed a notable stage-dependent pattern: the macronutrient associated with higher resistance often differed between development and adulthood. Larval-C extended HR (Mid F: P = 0.048, Supp. Table-ST4c; Late M: P = 0.0004, Supp. Table-ST4h) and OSR (Early F: P = 0.03, Supp. Table-ST3a; Mid F: P = 6.6 × 10⁻¹⁰, Supp. Table-ST3b; Mid M: P = 0.003, Supp. Table-ST3e; Figure 3A), whereas adult-P most consistently extended these traits. Conversely, larval-P extended SR in early-age females (P = 7.9 × 10⁻⁴; Figure 3A, Supp. Table-ST2a), whereas adult-C extended SR in early– and mid-age females. Thus, where larval and adult diet effects could be interpreted separately, developmental and adult nutrition did not act in the same macronutrient direction: the diet associated with higher resistance during development was often not the diet associated with higher resistance during adulthood.

Beyond these main effects, larval × adult interactions showed that some dietary responses were combination-specific rather than attributable to either dietary stage alone. This was most evident for DR and CR. DR was diet-sensitive in most sex–age strata but lacked a recurrent main-effect direction, with significant larval × adult interactions in early– and mid-age panels of both sexes (Early F: P = 0.02, Supp. Table-ST6a-b; Mid F: P < 2× 10⁻¹⁶, Supp. Table-ST3c-d,; Early M: P = 3 × 10⁻¹³, Supp. Table-ST3f-g; Mid M: P = 0.002 Supp. Table-ST3h-i; Figure 3A, S5). At late age, DR responses became independently interpretable: larval effects in females and both larval and adult effects in males. CR was similarly regime-specific. Apart from an independently interpretable larval-C effect in early-age (P = 0.03, Supp. Table-ST5a) and adult-C effect in mid-age females (P = 1.4 × 10⁻¹¹, Supp. Table-ST5b; Figure 3A), CR responses depended on the specific larval–adult combination (Supp. Table-ST5d,e,f, Figure 3A); notably, late-age CR converged on the same regime-level pattern in both sexes, with PC (larval-P/adult-C) showing the highest resistance and PP (larval-P/adult-P) the lowest(Supp. Table-ST5c, ST5g,h).

Together, these trait-level models show that dietary history modifies resistance selectively rather than shifting the covariance architecture uniformly. Adult diet generated the clearest directional signatures, developmental diet often acted in opposite macronutrient directions where independently detectable, and DR and CR were strongly shaped by larval–adult combinations. These analyses identify where diet acts trait by trait; we next asked whether those diet-induced shifts are coordinated across traits and whether they extend to fitness-associated outcomes.

### Adult-diet responses show a dietary-effect architecture across resistance and fitness-associated traits

Subsequently, we asked whether diet-induced responses were directionally concordant or opposed across resistance traits, and how these patterns aligned with fitness-associated outcomes. The dietary-effect index (DEI) summarizes whether the dietary response of one trait aligns directionally with that of another—a distinct question from covariance analyses, which ask whether traits vary together across conditions (Methods). We applied DEI separately to larval– and adult-diet contrasts and compared resistance responses against previously published longevity and fertility reference responses from the same dietary framework, as detailed in Methods.

Larval-diet effects produced relatively few significant pairwise DEIs (Table ST14 a, b, c, supplementary). Significant associations were sparse, occurring mainly in early-life females, rare at mid-life, and not at the later-life point in either sex. Thus, although developmental diet affected individual resistance traits, directional alignment between resistance and fitness-associated responses was not consistent across the examined strata.

Adult-diet effects, by contrast, showed more recurrent patterns within the dietary-effect architecture (Table 1; full matrix in supplementary Table ST15). HR, OSR, longevity, and fertility showed recurrent positive directional alignment: wherever significant, pair-wise DEIs for HR–OSR, HR–longevity, HR–fertility, OSR–longevity, and OSR–fertility were positive across the relevant comparisons. Thus, adult-diet responses of HR and OSR were directionally concordant with the corresponding longevity and fertility responses in the relevant strata.

**Table 1.**
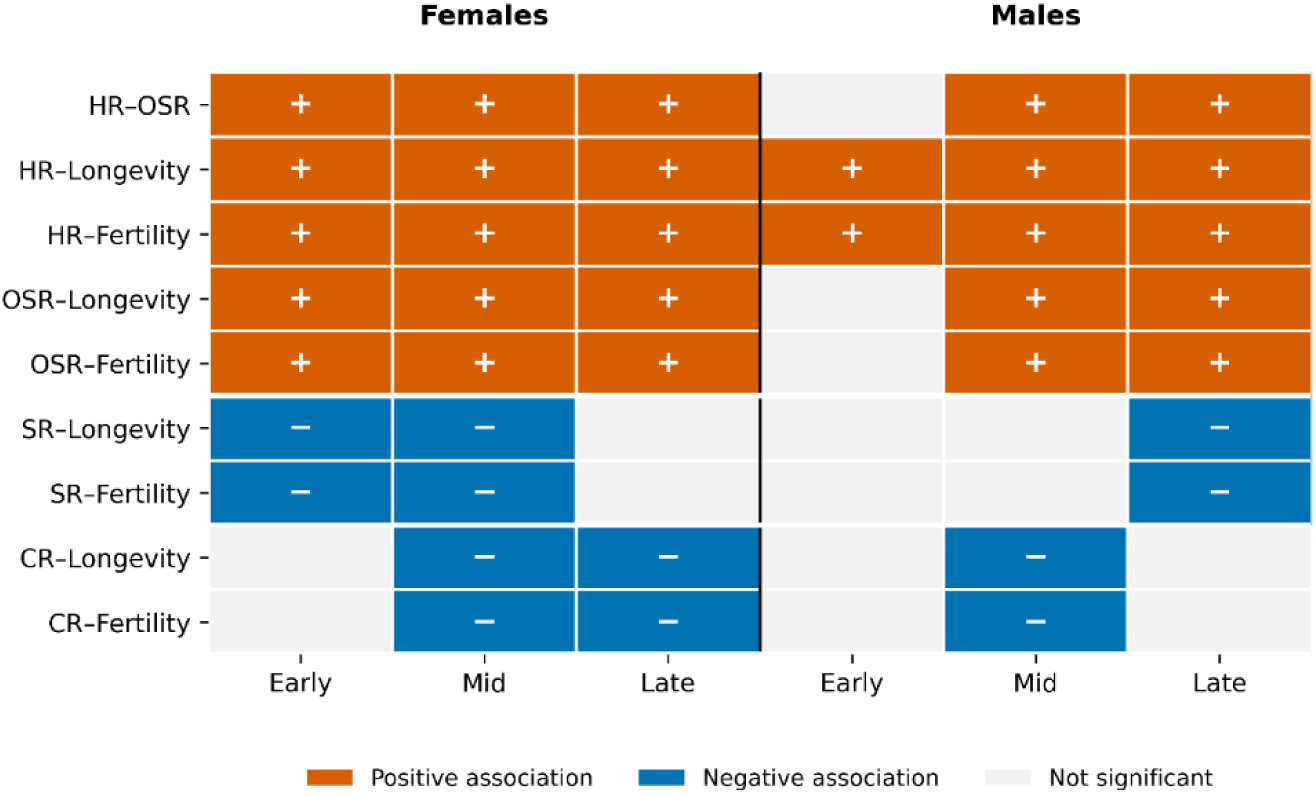
Adult-diet effect architecture linking stress-resistance traits with longevity and fertility across sex and age. Cells show adult-diet effect associations between trait pairs within each sex × age stratum. Orange indicates significant positive associations, blue indicates significant negative associations, and grey indicates non-significant associations. Plus and minus symbols denote association direction. HR, heat resistance; OSR, osmotic stress resistance; SR, starvation resistance; CR, cold resistance.

|  | Females |  |  | Males |  |  |
| --- | --- | --- | --- | --- | --- | --- |
|  | Early | Mid | Late | Early | Mid | Late |
| HR-OSR | + | + | + |  | + | + |
| HR-Longevity | + | + | + | + | + | + |
| HR-Fertility | + | + | + | + | + | + |
| OSR-Longevity | + | + | + |  | + | + |
| OSR-Fertility | + | + | + |  | + | + |
| SR-Longevity | - | - |  |  |  | - |
| SR-Fertility | - | - |  |  |  | - |
| CR-Longevity |  | - | - |  | - |  |
| CR-Fertility |  | - | - |  | - |  |

■ Positive association ■ Negative association ■ Not significant

SR showed the clearest opposing dietary-response pattern relative to the HR–OSR component. Wherever significant, SR–HR and SR–OSR DEIs were negative (Mid F and Late M; Supplementary Table ST15), indicating that adult diet shifted SR in the opposite direction from HR and OSR in these strata. SR was likewise negatively aligned with longevity and fertility in Early and Mid females and Late males (Table 1). CR showed a related but more context-dependent pattern, with negative dietary-response associations with HR and/or OSR and with longevity and fertility across selected mid– and late-life strata. Thus, SR in particular showed a dietary-response pattern distinct from the HR–OSR component and the corresponding fitness-associated responses.

Together, these results show that the covariance and dietary-effect architectures only partly correspond. HR and OSR formed part of the tightly correlated covariance core and also showed recurrent positive dietary-response alignment with longevity and reproductive output. SR, in contrast, remained peripheral to the covariance core yet, where significant, responded to adult diet in the opposite direction from HR/OSR and from the corresponding fitness-associated responses. CR similarly showed opposing dietary responses in several within-stratum comparisons. Thus, covariance among resistance traits did not reliably predict how those traits responded to nutritional perturbation. We next asked whether these contrasting adult-diet responses were accompanied by distinct whole-body physiological states.

### Whole-body triacylglyceride and protein content accompany contrasting adult-diet resistance signatures

We quantified whole-body triacylglyceride (TAG) and total protein across age to ask whether the adult-diet resistance signatures identified above were accompanied by divergent whole-body biochemical states. Because larval and adult diet effects were most clearly distinguishable in females, TAG and protein assays were focused on females to test whether these resistance signatures were accompanied by corresponding whole-body biochemical differences. Adult-C diet elevated TAG at all three ages (Early F: adult P = 0.0001; Late F: adult P = 3 × 10⁻⁸, Supp. Table-ST8a, ST8d), with Mid F showing a significant larval × adult interaction (P = 0.0179, Supp. Table-ST8b, ST8c) in which adult-C regimes consistently exhibited elevated TAG (Figure 4A–C). Conversely, P diet elevated whole-body protein, with the signature becoming increasingly adult-diet dominated across age: at Early F, any P exposure was sufficient to elevate protein (interaction P = 1.68 × 10^−04^, Supp. Table-ST9a,ST9b, Figure 4D), whereas by Mid F and Late F adult-P was the dominant driver (Mid F interaction P = 0.010; Late F interaction P = 0.421; Figure 4D–F, Supp. Table-ST9c, ST9d, ST9e, ST9f). Dry body weight broadly paralleled protein, with adult-P regimes showing higher dry weight at Mid F, Late F, and Mid M (Supp. Table ST7c, ST7d, ST7g, Figure S7A-C).

**Figure 4.**
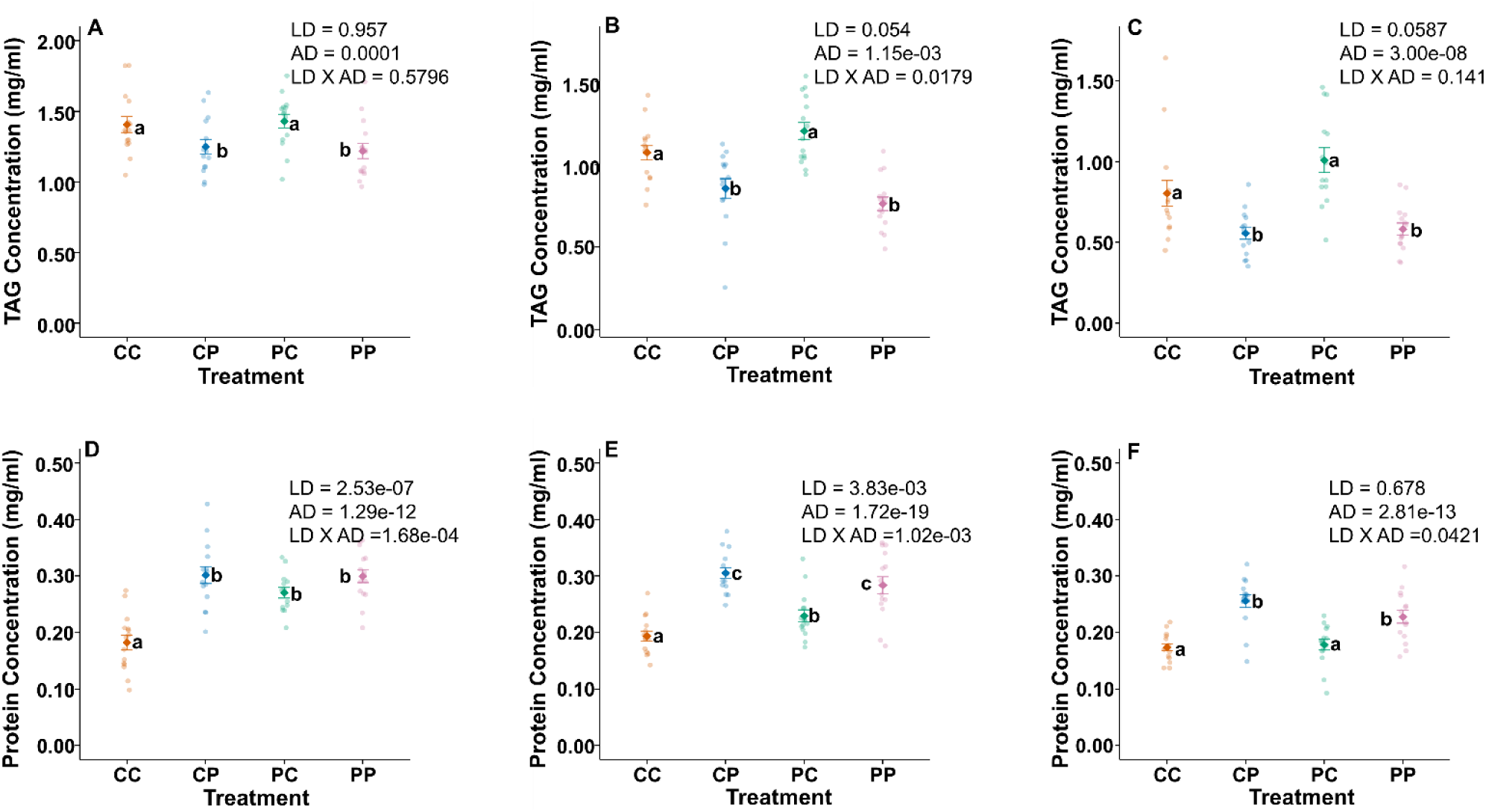
Adult diet generates contrasting TAG– and protein-associated physiological states in females. **A–C**, Whole-body triacylglyceride (TAG) concentration across larval × adult diet regimes in early (**A**), mid (**B**) and late (**C**) adult females. **D–F,** Whole-body protein concentration across the same diet regimes and ages. Coloured circles show biological replicates, and group means with error bars are shown in corresponding darker colours. Diet regimes are denoted by two letters, where the first letter indicates larval diet and the second indicates adult diet: CC, carbohydrate-biased larval and adult diet; CP, carbohydrate-biased larval and protein-biased adult diet; PC, protein-biased larval and carbohydrate-biased adult diet; PP, protein-biased larval and adult diet. P values indicate model-based tests of larval diet (LD), adult diet (AD) and their interaction (LD × AD). Different letters indicate significant statistical differences among diet regimes. Adult carbohydrate-biased diet was associated with elevated TAG, whereas adult protein-biased diet was associated with elevated protein concentration, indicating distinct adult-diet-dependent physiological states.

These biochemical patterns aligned directionally with the adult-diet resistance signatures. The adult diet that elevated TAG was associated with higher SR in female early– and mid-age strata, whereas the adult diet that elevated protein and dry weight preferentially raised HR and OSR. Thus, the contrasting adult-diet signatures corresponded to distinct whole-body biochemical states: an adult-C/TAG/SR-associated state and an adult-P/protein/HR–OSR-associated state. Greater body weight of adult-P flies (Fig. S7) alone, however, did not translate into higher starvation resistance: under starvation, adult-P flies lost a larger fraction of dry body weight at early and mid age (Early F: adult P = 7 × 10⁻⁴, Supp. Table ST10a; Mid F: adult P = 1.4 × 10⁻¹⁰, Supp. Table ST10b; Mid M: adult P = 2 × 10⁻⁴, Supp. Table ST10e; Fig. S8A-C), and this greater weight loss coincided with the lower SR of adult-P flies in female early– and mid-age strata.

Thus, within females, biochemical state and resistance phenotype showed consistent directional alignment. Whether these signatures also occur in males remains unresolved. These biochemical states accompany the dietary-effect signatures but do not explain what the tightest covariance core represents physiologically. Because DR, OSR, and HR formed the tightest covariance core, and because DR and OSR are directly linked to water balance, we then asked whether this core could be reduced to baseline or stress-associated water content.

### The integrated DR–OSR–HR resistance core extends beyond bulk hydration states

To test whether the tight DR–OSR–HR covariance core reflected shared water-balance physiology, we examined water-content traits alongside resistance traits in sex-stratified Pearson correlation matrices. Correlations were calculated across age × diet-regime combinations using estimated DR, OSR, and HR values together with baseline water content (WaterC) and water content measured at 10% mortality under osmotic or desiccation stress (WaterOSR and WaterDR; Figure 5; full water-content panels in Figs. S9-11).

**Figure 5.**
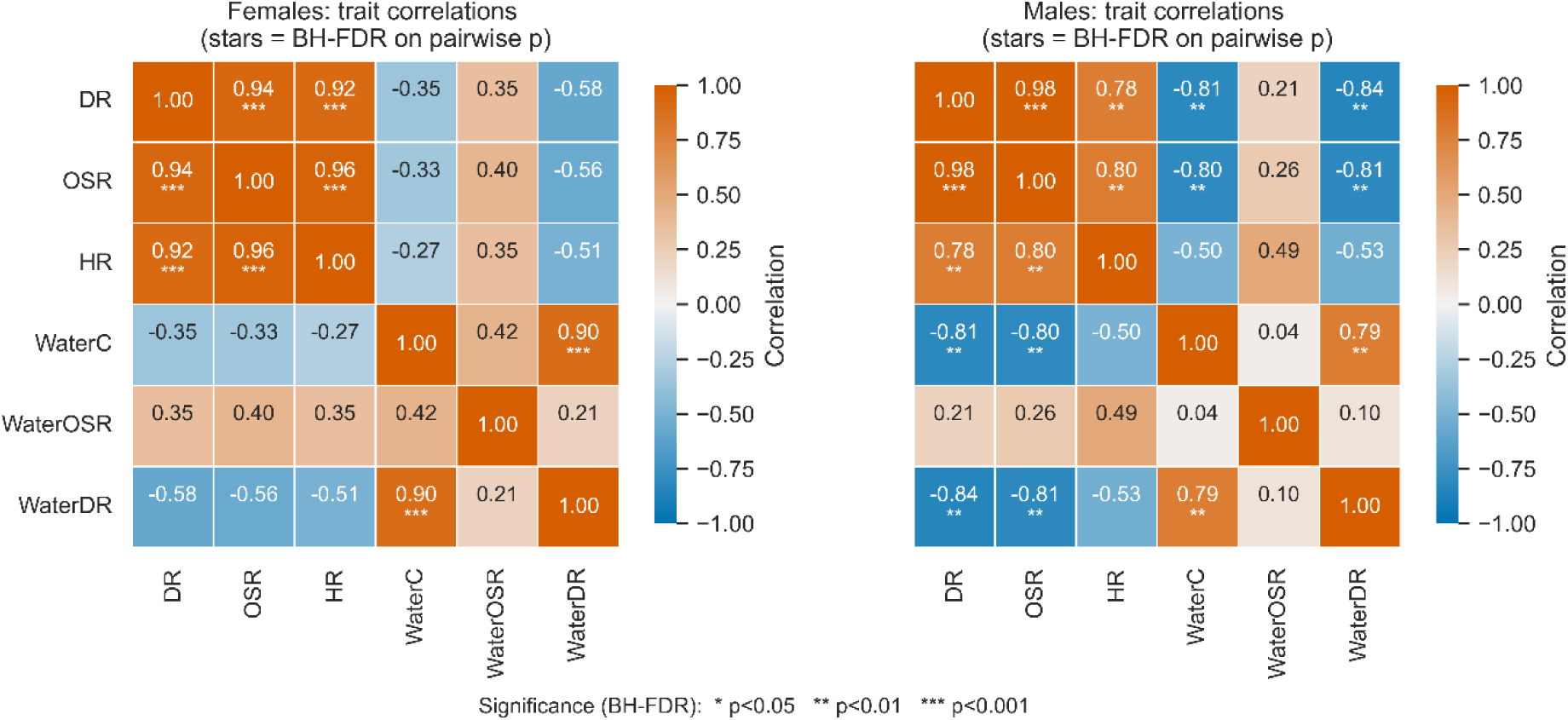
The integrated DR–OSR–HR resistance core is not explained by bulk water content. Sex-specific Pearson correlation matrices among desiccation resistance (DR), osmotic stress resistance (OSR), heat resistance (HR), baseline water content (WaterC), water content under osmotic stress (WaterOSR) and water content under desiccation stress (WaterDR) in females (left) and males (right). Correlations were calculated across condition-level means. Cell values indicate correlation coefficients, with colours showing the direction and magnitude of association. Asterisks denote BH–FDR-adjusted significance levels based on pairwise correlation: * *q* < 0.05, ** *q* < 0.01, *** *q* < 0.001. DR, OSR and HR remained strongly positively correlated in both sexes, whereas bulk water-content traits formed a partially distinct physiological structure.

Including water-content traits did not weaken the DR–OSR–HR core: DR, OSR, and HR remained strongly intercorrelated in both females (r = 0.94–0.96) and males (r = 0.84–0.98; Figure 5). By contrast, baseline water content did not positively align with this core. WaterC was weakly negatively associated with DR, OSR, and HR in females and more strongly negatively associated in males, including WaterC–DR (r = −0.81) and WaterC–OSR (r = −0.80). WaterDR showed a similar negative pattern, again strongest in males. WaterOSR was the only water-content trait that showed positive associations with DR, OSR, and HR (females: r = 0.34–0.41; males: r = 0.21–0.42), but these associations were substantially weaker than the internal DR–OSR–HR correlations and did not define the same integrated structure.

Water-content traits instead formed a partially distinct physiological structure (Figure 5). WaterC and WaterDR were strongly correlated in both females (r = 0.90) and males (r = 0.79), indicating that water content after desiccation retained much of the baseline water-content structure. Yet this water-content structure was decoupled from the DR–OSR–HR core and, especially in males, negatively associated with resistance. Thus, the tightest resistance core was not explained by higher starting hydration or higher endpoint water content after stress.

Together, these results show that the DR–OSR–HR core is not a passive reflection of bulk water content. Instead, it represents a coordinated physiological state that persists independently of baseline and endpoint hydration measures, and in males is opposed to the major WaterC–WaterDR hydration structure.

## Discussion

Across the four traits included in the covariance analysis, stress resistance was dominated by a strong age-associated multivariate axis but was not uniform across traits. The covariance architecture was broadly similar between sexes: DR, OSR, and HR formed a tightly correlated core, whereas SR remained comparatively peripheral. This supports an intermediate position between generalized stress resistance and stress-specific physiology: resistance is neither a single uniform phenotype nor a collection of fully independent traits ^5,15,28^. Instead, the strongest shared component resides in the DR–OSR–HR core. Nutritional perturbation revealed a complementary level of organization, as patterns of dietary response did not simply recapitulate this covariance structure.

Although immediate effects of adult diet on adult stress resistance are expected, the important result was the form of this response. Adult nutrition did not simply shift resistance globally upward or downward; it produced recurrent but contrasting trait-specific responses, with adult-P generally favouring HR and OSR, whereas adult-C favoured SR in the relevant female strata. Developmental diet was not irrelevant, but its effects were more trait-, age-, and combination-specific: larval diet affected selected traits, DR and CR were strongly shaped by larval × adult combinations, and the beneficial macronutrient sometimes differed between developmental and adult stages. The persistence of some larval-diet effects into mid and late adulthood demonstrates long-lasting developmental carry-over, although whether this reflects persistent resource differences, developmental programming, or other physiological changes remains unresolved. Previous work has shown that larval diet can shape adult stress tolerance and that adult diet need not compensate for developmental nutritional history ^29,30^. Our results extend this view by showing that for traits in which larval and adult effects could be interpreted separately, the macronutrient associated with higher resistance sometimes differed between developmental and adult stages. Thus, the data point to persistent but trait-specific developmental carry-over alongside more recurrent adult-diet effects. The prominent influence of adult diet is particularly important because it shows that age-dependent stress performance remains nutritionally modifiable after development. Rather than uniformly improving resistance, adult macronutrient balance favoured different resistance traits, cautioning against treating improvement in any single stress phenotype as evidence of generalized physiological improvement.

Previous studies have likewise found that relationships among stress-resistance traits are only partly shared and depend on the context in which they are examined. Artificial-selection experiments in *Drosophila* have produced correlated responses among several stress traits, but these responses are neither universal nor reciprocal. For example, starvation resistance increased as a correlated response to selection for several other stress-resistance traits, whereas selection for starvation resistance itself did not produce broad cross-resistance ^5^. Relationships among desiccation and thermal resistance are similarly variable: desiccation selection has produced correlated heat– and cold-resistance responses in some experiments, whereas others found little increase in basal or rapid cold tolerance, apart from modest improvement in chill-coma recovery ^5,31^. Genetic studies provide a complementary perspective. QTL mapping has identified partially overlapping genomic regions affecting desiccation, cold, and heat resistance, although many effects are trait– or sex-specific and shared loci can have opposing effects across stress phenotypes^32–34^. Thus, both selection and genetic-mapping studies support partially shared rather than unitary stress-resistance architecture. Our results extend this perspective by separating two aspects of this structure within the same nutritional experiment: DR, OSR and HR formed a strong covariance core across conditions, with SR remaining peripheral, yet the responses of these traits to nutritional perturbation did not simply reproduce that covariance pattern. Where significant, adult-diet responses of SR were opposed to those of HR and OSR, despite the absence of antagonistic covariance among these traits. Thus, covariance identifies which resistance traits tend to vary together across conditions, but does not necessarily predict how those traits respond when the nutritional environment is altered.

This distinction also extends to fitness-associated outcomes in our results. Adult-diet responses of HR and OSR were recurrently aligned with longevity and reproductive output, whereas SR—and, in several within-stratum comparisons, CR—responded in the opposite direction from these reference responses. The relationship between stress resistance and longevity was therefore trait-specific and nutritionally contingent rather than a general property of stress resistance, consistent with experimental evidence that stress resistance and lifespan can be dissociated ^6^. This pattern should not be interpreted as evidence of a genetic or evolutionary trade-off, because the present analysis compares directional responses to diet rather than causal allocation constraints. It is nevertheless consistent with acquisition– allocation theory, under which positive covariance can arise when conditions differ in resource acquisition, while perturbing the resource environment can reveal contrasting responses among allocation-dependent traits ^8^. In this sense, predominantly positive covariance among the resistance traits included in the covariance analysis and opposing dietary responses among particular traits are not contradictory; they capture different aspects of phenotypic response.

These results also complicate a simple protein-mediated opposition between lifespan and reproduction. In nutritional-geometry studies, lifespan and reproduction often map to different macronutrient optima, with reproductive traits generally favoured by higher protein availability and lifespan often favoured by lower-protein, carbohydrate-biased diets ^9,35^. Here, adult-diet responses of HR and OSR were positively aligned with both longevity and reproductive output, consistent with recent work in this outbred *Drosophila* system showing that protein-biased diets can increase both fecundity and lifespan under isocaloric macronutrient manipulation ^13^. Thus, macronutrient effects on ageing-related performance may vary in a trait-specific manner rather than following a single lifespan–reproduction axis. Nutrient-sensing pathways such as IIS/TOR provide plausible mechanistic candidates for these patterns, because they link amino-acid and carbohydrate availability to growth, reproduction, storage, proteostasis, and somatic maintenance. However, because pathway activity was not measured directly, our results identify candidate physiological states and phenotypic relationships rather than causal molecular mechanisms.

The physiological measurements provide context for these dietary-response patterns rather than a causal explanation. Adult-C diet produced a TAG-rich state aligned with SR, consistent with prior evidence that lipid reserves contribute to starvation resistance in *Drosophila* ^22,36^. Adult-P produced higher whole-body protein and dry weight alongside higher HR and OSR, identifying a distinct biochemical state associated with these resistance responses. This alignment is physiologically plausible because heat and osmotic stress both require active maintenance of cellular homeostasis: heat tolerance depends partly on proteostasis and heat-shock responses, whereas osmotic tolerance depends on ion and fluid regulation, including epithelial transport systems such as the Malpighian tubules ^37–40^. Greater body mass alone, however, did not translate into higher starvation resistance, because adult-P flies lost a larger fraction of dry body mass under starvation and showed lower SR in the relevant female strata. This pattern is consistent with the idea that starvation resistance depends not only on the amount of stored material, but also on the form of storage and the rate at which reserves are mobilized under deprivation. Lower TAG availability in adult-P females may contribute to greater reliance on non-lipid reserves during starvation, although this remains to be tested directly. These results suggest that the adult-C/TAG/SR and adult-P/protein/HR–OSR signatures reflect distinct nutritional states rather than a simple effect of overall body size.

The DR–OSR–HR covariance core was not reducible to baseline or endpoint water content. Bulk hydration formed a partially distinct physiological structure and, especially in males, was negatively associated with resistance. This argues against a simple model in which higher starting water content or residual water after stress explains the covariance core. Instead, the DR–OSR–HR core may reflect shared physiological processes involving proteostasis, osmotic and ion regulation, cuticular permeability, water-loss control, or shared stress-response pathways ^39,41–43^. The positive dietary-response alignment of HR and OSR with longevity and reproductive output further suggests that these components of the core may be associated with physiological states relevant to ageing performance, although the causal basis of this relationship remains to be tested. Thus, the DR–OSR–HR covariance core cannot be explained simply by greater bulk hydration, while its underlying physiological basis remains to be resolved.

Several limitations should be considered. The study used two contrasting P:C diets rather than a broader nutritional gradient; therefore, the observed relationships should be interpreted within this dietary contrast rather than as defining the full nutritional response surface. Because P:C ratio was manipulated by varying yeast and sugar, the diets also differed in other yeast-derived nutrients, including sterols, which can influence lifespan responses to dietary manipulation in *Drosophila* ^44^. Moreover, equal caloric density does not ensure equal caloric intake, which was not measured. This is also relevant to the high-salt assay, where variation in feeding could influence NaCl ingestion. Although starvation and high-salt resistance showed distinct dietary-response profiles, reduced feeding under high-salt conditions may still contribute to survival. TAG and protein were measured only in females, limiting inference about whether the same biochemical alignments explain male resistance architecture. Water content was measured at condition-level endpoints rather than as dynamic water-loss rates. As cold-challenge duration differed among sex × life-course strata, CR was excluded from cross-stratum covariance analyses and retained only for trait-specific and dietary-response comparisons within strata, where challenge duration was identical across dietary treatments. Because sampling was scheduled by time post egg collection rather than post eclosion, diet-dependent differences in developmental time mean that larval-diet groups may differ in time since eclosion and cumulative adult-diet exposure. Larval-diet effects should therefore be interpreted in the context of these fixed post-egg-collection life-course points rather than as effects measured at matched post-eclosion ages. The links to longevity and fertility describe aligned dietary responses across matched datasets, not individual-level physiological coupling. Future work combining male and female biochemistry, lipidomics, osmolyte profiling, cuticular hydrocarbon and permeability assays, metabolic-rate measurements, and nutrient-sensing perturbations will be needed to identify the mechanisms linking adult diet, stress resistance, and fitness-associated outcomes. These experiments could test whether the adult-P/protein/HR–OSR and adult-C/TAG/SR signatures reflect differences in proteostasis, osmoregulation, storage, or other aspects of somatic physiology. Finally, although the use of an outbred *Drosophila* population captures substantial standing genetic variation, genetic background should be considered when generalising these findings, as genotype-by-diet interactions can generate distinct phenotypic responses ^45^.

Together, these findings show that stress resistance across the adult life course is structured but not uniform across traits. Among the traits included in the covariance analysis, DR, OSR, and HR formed a shared covariance core in both sexes, whereas SR remained comparatively peripheral, with life-course progression dominating the multivariate pattern. Nutritional perturbation revealed a complementary but non-equivalent pattern: adult diet produced recurrent contrasting responses among resistance traits, while developmental diet left persistent but more context-dependent effects. HR and OSR showed dietary responses aligned with longevity and reproductive output, whereas SR—and CR in several within-stratum comparisons—showed opposing responses, accompanied by distinct whole-body physiological states. Thus, the covariance structure among stress-resistance traits was not sufficient to predict their responses to nutritional perturbation, revealing a distinction between how traits covary across conditions and how they respond when nutrition is altered. More broadly, stress resistance emerges as a differentiated component of ageing physiology whose relationships depend on nutritional history, life-course stage, and the particular stress phenotype considered.

## Methods

### Experimental population and maintenance

We used BP_4_, one of five replicate outbred populations derived from a common Baseline Population (BP)^13^. The BP was founded by equally mixing the MB_1_–MB_4_ populations, whose ancestry is described elsewhere (Sarangi et al. 2016; Shenoi et al. 2016), and was maintained for 28 generations before being split into BP_1_–BP_5_. These populations have since been maintained independently under identical conditions with 25 °C, 60–70% relative humidity, and constant light on corn–sugar–yeast (CSY) medium prepared following an established recipe ^46^, with a protein-to-carbohydrate ratio of 0.4 by mass calculated from ingredient composition, during both larval and adult stages under routine maintenance ^13^. Preadult development occurred in fly bottles (Laxbro® FLBT-20) containing 50 mL food at 300-350 eggs per bottle (for details, see supplementary Text S2). Adults emerging from seven bottles were pooled into Plexiglas cages (25 × 20 × 15 cm; L × W × H), containing the same medium, with no live-yeast supplement, giving a census of approximately 2000 adults per generation. An egg-laying surface was provided overnight on day 13 post egg collection, and eggs were collected on day 14 to found the next generation, giving a 14-day discrete-generation cycle. BP_4_ had been maintained on this regime for 42 generations since the split when the present experiments began. This is the baseline maintenance regime; experimental diets are described in the next section.

### Dietary paradigm and experimental design

We implemented an isocaloric dietary manipulation following established protocol ^24^, using two protein-to-carbohydrate (P:C) ratios: 0.7 (protein-biased; henceforth, P diet) and 0.25 (carbohydrate-biased; henceforth, C diet). These ratios were selected based on prior experiments showing that both isocaloric diets produced preadult viability and adult body size comparable to the intermediate baseline diet, indicating that neither represented a malnutritive extreme ^13^.

Diet was manipulated independently during larval development and adulthood in a full-factorial larval diet × adult diet design. Eggs from the baseline population were collected at approximately 150 eggs per 50 mL food in plastic fly bottles containing either the P or C larval diet. At the late pupal stage, identified by pupal darkening, the detachable food-containing base of each bottle was replaced with one containing the assigned adult diet, ensuring exposure to the adult diet immediately upon eclosion. Each larval treatment was thus split between P and C adult diets, generating four regimes: PP, PC, CP, and CC, where the first letter denotes larval diet and the second adult diet. On day 12 post egg collection, all eclosed adults from each regime were transferred to a dedicated mixed-sex Plexiglas cage and maintained at densities not exceeding approximately 2,000 flies, with fresh respective adult diet provided in Petri dishes every other day.

All stress-resistance and physiological assays were conducted cross-sectionally on days 12, 22 and 32 post egg collection, representing early-, mid-, and later-life points across the adult life course. Because developmental diet influences developmental time, these fixed post-egg-collection points do not correspond to precisely matched post-eclosion ages across larval-diet treatments. These ages were chosen using population-specific survival data to sample adult function across ageing while limiting differential survivorship among dietary treatments at the latest age ^13,24,26^.

Each assay used separate flies sampled from the corresponding treatment populations; therefore, individual flies were not measured repeatedly across traits. Multivariate analyses described below consequently quantify covariance among trait values across experimental conditions rather than within-individual covariance.

### Stress-resistance assays

#### Starvation resistance

Flies were aspirated directly from treatment cages and separated by sex without CO₂ anaesthesia. Individual flies were placed in glass activity-monitor tubes (80 mm length × 5 mm diameter) and loaded into DAM2 Drosophila Activity Monitors. One end of each tube was closed with cotton and the other with a 0.2 mL PCR tube containing 1% agar as a water source. Agar tubes were replaced every 48 h to limit microbial growth.

Activity was recorded at 5-min intervals. Starvation resistance was defined as the interval from assay initiation to the last recorded activity. A total of 954 flies were assayed across dietary regimes, sexes, and ages.

#### Desiccation resistance

Flies were separated by sex under brief, mild CO₂ anaesthesia. Groups of eight same-sex flies were placed in empty glass vials without food or water, with 10 replicate assay vials per treatment combination. Survival was scored every 2 h until all flies had died. Desiccation resistance was defined as time from assay initiation to death. In total, 1,918 flies were assayed across dietary regimes, sexes, and ages.

#### Heat resistance

Acute heat resistance was measured individually using DAM2 Activity Monitors. Flies were aspirated from treatment cages and placed individually in activity-monitor tubes containing their corresponding adult diet at one end. The food end was sealed with Parafilm and the opposite end with cotton.

Assays were conducted at 35 °C under constant light. Activity was recorded at 30-s intervals, and heat resistance was defined as the interval from assay initiation to the last recorded activity, used as a proxy for death. Approximately 32 flies were assayed per diet regime, sex, and age, giving a total sample size of 767 flies.

#### Cold resistance

Cold resistance was quantified as survival following prolonged exposure to 1 °C. Flies were aspirated from the corresponding treatment cages and placed individually in empty DAM tubes sealed with cotton at both ends. Tubes were enclosed in small resealable plastic bags and transferred to 1 °C.

Because baseline cold susceptibility differed with sex and age, exposure durations were standardized separately for each sex × age group but held identical across the four dietary regimes within that group. Females were exposed for 36, 28, and 24 h at early, mid, and late age, respectively; males were exposed for 24, 20, and 16 h, respectively. Following exposure, flies were returned to room temperature and allowed to recover for 1 h before survival was scored. Approximately 30–35 flies were assayed per dietary regime within each sex × age group, giving a total sample size of 790 flies.

#### Osmotic stress resistance

Osmotic-stress resistance was measured under high-salt dietary conditions. Males and females were separated under brief, mild CO₂ anaesthesia, and groups of five same-sex flies were placed in vials containing the corresponding adult diet supplemented with 4% NaCl. Ten replicate assay vials were used per treatment combination. Survival was scored every 2 h until death, and alive flies were transferred to fresh food vials every alternate day until all flies in a vial had died. Osmotic-stress resistance was defined as time from assay initiation to death. In total, we assayed 1200 flies across dietary regimes, sexes, and ages. To assess whether the composition of the assay food contributed to the observed high-salt resistance differences, a targeted validation assay was conducted at day 22 post egg collection. Flies from all prior dietary treatments were challenged on a common P:C = 0.4 food containing 4% NaCl, thereby standardising the basal assay-food composition and same NaCl concentration across treatments while preserving their distinct nutritional histories. All other assay conditions were identical to the original OSR assay.

#### Longevity and Fertility

Previously published longevity data^14^ and fertility data^27^ were obtained from the same outbred population under the same full-factorial larval × adult dietary design used here. Fertility was assayed on days 12, 22, and 32 post egg collection as the number of viable progeny produced during a standardized 24-h reproductive assay^27^. Thus, the fertility measure used here represents age-specific reproductive output rather than cumulative or lifetime reproductive success.

### Whole-body physiological measurements

#### Triacylglyceride content

Whole-body triacylglycerides (TAG) was quantified in females using a colorimetric assay kit (Sigma, TR0100) with minor modifications. Five flies constituted one biological replicate, with 15 biological replicates per treatment. Flies were homogenized in 150 μL PBST (PBS containing 0.005% Tween-20) using a bead beater at 50 Hz for three 30-s cycles separated by 10-s intervals. Homogenates were centrifuged at 3,000 rpm for 6 min to pellet debris.

For each biological replicate, 2 μL supernatant was assayed in duplicate. Endogenous glycerol was quantified after addition of 160 μL Free Glycerol Reagent and incubation at 37°C for 5 min. Plates were centrifuged at 3,000 rpm for 5 min at 4 °C to remove bubbles before absorbance was measured at 540 nm. Total TAG was then measured after addition of 40 μL Triglyceride Reagent, followed by a further 5-min incubation at 37 °C and absorbance measurement at 540 nm. A glycerol standard curve was included on each plate.

Technical duplicates were averaged before statistical analysis, and the biological replicate was treated as the unit of analysis. TAG concentrations were calculated from the corresponding standard curve and expressed as TAG concentration.

#### Total protein content

Whole-body protein was quantified in females using the bicinchoninic acid assay (Thermo Fisher Scientific, catalogue no. 23225). Five flies constituted one biological replicate, with 15 biological replicates per treatment. Flies were frozen at −20 °C for 20 min and homogenized in 200 μL RIPA buffer containing protease inhibitor (Merck, catalogue no. 11836153001). Homogenates were centrifuged at 10,000 rpm for 3 min at 4 °C.

A 10 μL aliquot of supernatant was diluted to fall within the linear range of the assay, and 25 μL diluted lysate was combined with 200 μL BCA working reagent prepared at a 50:1 ratio of reagents A-to-B. Plates were incubated at 37 °C for 30 min and absorbance was measured at 562 nm. Protein concentration was calculated from a bovine serum albumin standard curve included on each plate. Two technical replicates were averaged before analysis.

### Dry body weight and starvation-associated weight loss

Dry body weight was quantified in age-matched control and starvation-exposed flies. Groups of 10 flies of the same sex, age, and treatment were placed in pre-labelled 2 mL microcentrifuge tubes, dried at 60 °C for 72 h, and weighed to the nearest 0.1 mg using a high-precision analytical balance (Sartorius BCE223i-1S). Ten replicate tubes were processed per treatment.

For the starvation-exposed group, flies were subjected to the starvation protocol described above and sampled when approximately 10% mortality had occurred. Corresponding control flies were maintained on their respective adult diets over the same interval. Dry weight was measured identically in both groups. Absolute mass loss was calculated as control dry mass minus post-starvation dry weight, and proportional weight loss as:

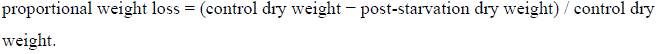

### Water content before and during desiccation and hyperosmotic stress

Whole-body water content was estimated from fresh and dry body weight. Flies were separated by sex using aspiration, euthanized under diethyl ether, and weighed immediately to the nearest 0.1 mg. Samples were then dried at 60 °C for 72 h and reweighed.

Absolute water content was calculated as fresh weight minus dry weight. For analyses based on proportional water content, water fraction was calculated as:

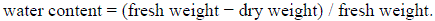

To estimate water content during desiccation and osmotic stress, separate groups of flies were exposed to the corresponding stress assay and collected when approximately 10% mortality had occurred. Fresh and dry weights were then measured using the same procedure. These measurements are referred to as baseline water content (WaterC), water content during desiccation stress (WaterDR), and water content during osmotic stress (WaterOSR). Ten replicates with 10 flies in each were processed per sex × age × dietary-regime combination.

## Statistical Analysis

### General analysis strategy

Analyses were conducted at two complementary levels. First, trait-specific models tested how larval diet, adult diet, age, and sex affected individual resistance and physiological traits. Second, condition-level multivariate analyses examined how resistance traits covaried across experimental conditions and whether dietary effects on different traits were directionally aligned.

Models were specified a priori according to the full-factorial experimental design and were not selected by stepwise simplification. Initial models included larval diet, adult diet, age, sex, and their interactions. Because dietary effects differed among sexes and ages, subsequent analyses were stratified by sex and age to estimate larval-diet effects, adult-diet effects and larval × adult diet interactions.

Where a larval × adult diet interaction was significant, the dietary response was interpreted as combination-specific and marginal larval– or adult-diet main effects were not interpreted independently. Where the interaction was non-significant, larval– and adult-diet main effects were interpreted separately. All tests were two-sided, with α = 0.05 unless otherwise stated.

### Trait-specific models

Time-to-event data for starvation, desiccation, heat, and osmotic-stress resistance were analysed using Cox proportional-hazards models. For desiccation and osmotic-stress assays, assay vial was included as a random intercept to account for the shared environment of flies housed within the same vial, using mixed-effects Cox models implemented in the *coxme* package. Starvation and heat resistance were measured in individually housed flies and were analysed without an assay-vial random effect. Cold resistance was analysed as a binary survival outcome using binomial generalized linear models with a logit link. TAG, protein, and dry body weight were analysed using Gaussian linear models. Proportional weight loss and proportional water-content variables were arcsine-square-root transformed before analysis and analysed using Gaussian linear models.

### Condition-level summaries for multivariate analyses

One condition-level summary was obtained for each resistance trait within every sex × age × dietary-regime combination. The four larval–adult dietary regimes and three ages produced 12 conditions per sex.

For starvation, desiccation, heat and osmotic-stress resistance, survival was summarized using restricted mean survival time (RMST). Kaplan–Meier curves were estimated separately for each sex × age × dietary-regime condition. Within each assay, a single common truncation time, *τ*, was applied to all conditions and was defined as the maximum observed follow-up time in the complete assay dataset. RMST was calculated as the area under the Kaplan–Meier survival curve from time zero to *τ* using the Python lifelines package.

Cold resistance used a distinct endpoint design. Flies were exposed to a sex– and life-course-specific cold challenge and scored as alive or dead after a subsequent 1-h recovery period.

Because challenge duration differed among sex × life-course strata, CR was not included in cross-stratum PCA, correlation-network, or integration analyses. CR was retained in trait-specific and dietary-effect analyses conducted within each stratum, where exposure duration was identical across dietary treatments.

The RMST values for SR, DR, OSR, and HR were assembled into a single covariance-analysis matrix containing 24 condition-level observations: 12 life-course × dietary-regime conditions for each sex. Before multivariate analysis, each trait was standardized across the complete 24-condition dataset to mean zero and unit variance.

*Principal component analysis*: Principal component analysis (PCA) was performed separately for females and males on the standardized condition-level matrix containing SR, DR, OSR, and HR. For each sex, the analysis therefore included 12 age × dietary-regime conditions. PCA was implemented in *Python* using *scikit-learn*, and the first two principal components were retained for visualization. PCA was used descriptively to identify the dominant dimensions of multivariate resistance variation and to visualize the distribution of age × dietary-regime conditions in multivariate trait space. To formally assess life-course structuring of the PCA, PC1 and PC2 scores were analysed separately within each sex using linear models with age and dietary regime as additive factors.

*Pairwise trait correlations and correlation networks*: The condition-level covariance architecture of stress resistance was characterized separately in females and males using Pearson correlations among SR, DR, OSR, and HR across the 12 age × dietary-regime conditions. Because the resistance traits were measured in separate flies, these correlations quantify correspondence among condition-level trait values rather than within-individual phenotypic correlations.

For each sex, P values were calculated for the six unique pairwise correlations and adjusted using the Benjamini–Hochberg false-discovery-rate procedure. Correlation networks contained all four traits as nodes and retained an edge only when the corresponding BH– FDR-adjusted P value was <0.05. Edge width was proportional to the absolute Pearson correlation coefficient, and edge colour distinguished positive from negative correlations.

## Dietary effect index

We compared whether pairs of traits responded to diet in the same or opposite directions, analysing each sex, age, and stage of dietary exposure separately, following previous literature ^24^. In short, for each trait, dietary response was quantified using the log response ratio, ln 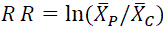. Positive values indicate higher trait values under the P diet, whereas negative values indicate higher values under the C diet. Sampling variance was estimated from the corresponding group means, standard deviations, and sample sizes. For each trait pair *i*, *j*, the dietary-effect index was calculated as *DEI_ij_* = ln *R R_i_*/ ln *R R_j_*, with uncertainty estimated using standard error propagation assuming independence between trait estimates. Statistical significance was assessed using two-sided z tests, as described previously. Positive DEI values indicate that both traits responded to diet in the same direction, whereas negative values indicate that one trait increased under the P diet while the other increased under the C diet. DEI therefore describes directional concordance or opposition between dietary responses and does not, by itself, imply functional coordination or mechanistic coupling. A negative DEI likewise does not show that improvement in one trait necessarily comes at the cost of the other.

For resistance–resistance comparisons, DEIs were calculated within each sex × age stratum. For comparisons with reproductive output, measurements from days 12, 22 and 32 post egg collection were matched to the corresponding resistance sampling points; because reproductive output was measured from male–female pairs, the same age-specific reproductive response served as the reference for male and female resistance traits. Longevity was a lifetime measure; therefore, the sex-specific dietary response in longevity was used as a common reference across the three resistance age points. These comparisons quantify directional alignment with the corresponding reproductive or lifetime longevity response rather than independent sex × age-specific fitness associations.

## Data availability

All data will be available on Dryad upon acceptance.

## Author contributions

Conceptualization: Devashish Kumar, Sudipta Tung

Design of the experiment: Devashish Kumar, Mohankumar Chandrakanth, Chetan S, Nishant Kumar, Chand Sura, Sudipta Tung

Data curation: Devashish Kumar, Mohankumar Chandrakanth, Chetan S, Nishant Kumar, Chand Sura, Sudipta Tung

Formal analysis: Devashish Kumar

Funding acquisition: Sudipta Tung

Supervision: Sudipta Tung

Visualisation: Devashish Kumar, Sudipta Tung

Writing – original draft: Devashish Kumar, Sudipta Tung

Writing – review & editing: Devashish Kumar, Mohankumar Chandrakanth, Chetan S, Sudipta Tung

## Acknowledgements

DK, CH, NK and CS acknowledge the support of the Research and Development Office, Ashoka University. MC acknowledges the support from the Department of Biotechnology, Govt. of India through Junior Research Fellowship. ST acknowledges the support of DBT/Wellcome Trust India Alliance Early Career Fellowship (#IA/E/18/1/504347) and Ashoka University. Authors thank Ruchitha BG, Itibaw Farooq, Jeevak Shravasti for experimental help, and Akshay Malwade for comments on the initial version of the manuscript. Authors thank Sushil, Shiva, Darshan, Vikas, Sahil, and Virender for logistics and maintenance support.

