## Supplementary for "Nutritional history affects stress-resistance architecture across ageing and sex"

for

**Supplementary Results:**

**Text S1:** **Permutation-based integration analysis.**

Overall integration among the resistance traits was quantified separately within each sex from the eigenvalue distribution of the complete, unthresholded Pearson correlation matrix. For a matrix with $p$ eigenvalues, $\lambda_{1}\geq\cdots\geq\lambda_{p}$, the integration index was calculated as

$$I=\frac{\mathrm{Var}\left( \lambda_{1} , \ldots, \lambda_{p} \right)}{p},$$

where $\mathrm{Var}$denotes the population variance of the eigenvalues. Because the eigenvalues of a $p\times p$correlation matrix sum to $p$, larger values of $I$indicate a more uneven eigenvalue spectrum and greater concentration of trait variation into fewer shared dimensions. The proportion of total standardized trait variance captured by the first eigenvalue, $\lambda_{1}/p$, was reported as a complementary descriptor. Pearson correlations, and therefore $I$, are unaffected by the centring and rescaling used to standardize the traits.

Statistical significance was assessed using $B=50,000$permutations. In each permutation, the 12 condition-level values were shuffled independently within each trait, preserving each trait’s marginal distribution while disrupting systematic correspondence among traits across experimental conditions. A new Pearson correlation matrix and integration index were calculated for each permuted dataset. The permutation P value was calculated as

$$P=\frac{b+1}{B+1},$$

where $b$ is the number of permuted indices greater than or equal to the observed value. Adding one to both the numerator and denominator provides a valid, non-zero Monte Carlo permutation P value for a finite set of random permutations (Phipson and Smyth 2010). Integration was calculated independently for females and males and tested against a separate null distribution for each sex. The two observed sex-specific indices were therefore compared descriptively, because the analysis did not directly test the between-sex difference.

**Text S2:** **Egg collection protocol**

To collect eggs from adults reared in plexiglass chambers, we utilised oviposition surfaces (cut plates) that provided vertical surfaces for egg-laying. The oviposition surface was of the same dietary composition as the respective adult foods. The flies were permitted to oviposit on these plates overnight, for roughly 14 hours. For egg collection, the plates were gently removed from each cage, and their vertical surfaces were flushed with a small volume of distilled water to facilitate egg removal. For every cage, we pre-prepared an autoclaved 2 mL microcentrifuge tube containing 750 µL of distilled water. Using a fine, dampened paintbrush (size 000), we carefully collected the eggs off the plate and pooled them into the corresponding tube until the accumulated egg volume reached the 500 µL calibration mark.

To remove residual food debris from the sample, we added an additional 750 µL of distilled water to each tube. The tubes were then gently inverted to break up any clumps to ensure thorough mixing. After allowing the eggs to settle, we carefully pipetted and discarded 750 µL of the supernatant, which contained the bulk of the suspended food particles. Finally, we dispensed a 50 µL aliquot of the egg suspension into culture bottles prefilled with 50 mL of fresh medium. Based on our prior standardisation, this volume consistently seeded each bottle with approximately 300–350 eggs. For experimental population egg collection, the number of eggs was kept at 150-200 by adding 750 µL of water after the washing step to avoid any crowding effect. The cultures were subsequently incubated at 25°C under continuous illumination and maintained at a relative humidity exceeding 60%.

**Supplementary Table ST1**. Cold exposure durations* used for the cold resistance assay

| Sex | Age (days) | Exposure duration (h) |
| --- | --- | --- |
| Male | 12 | 24 |
|  | 22 | 20 |
|  | 32 | 16 |
| Female | 12 | 36 |
|  | 22 | 28 |
|  | 32 | 24 |

* Exposure durations were selected based on prior survival data to achieve ~50% mortality and were applied identically across dietary regimes.

**Supplementary Table ST2:**

**ST2: Results of age and sex stratified analysis for Starvation resistance**

(model: coxph(Surv(Time, Status) ~ Larval_food*Adult_food, data = data))

**ST2a:** Females (Early age)

|  | Loglik | Chisq | Df | Pr(>\|Chi\|) |
| --- | --- | --- | --- | --- |
| NULL | -640.278 | NA | NA | NA |
| Larval_food | -634.267 | 12.02322 | 1 | 0.0005254 |
| Adult_food | -631.808 | 4.916367 | 1 | 0.0266034 |
| Larval_food: Adult_food | -631.297 | 1.023464 | 1 | 0.3116988 |

**ST2b:** Females (Mid-age)

|  | Loglik | Chisq | Df | Pr(>\|Chi\|) |
| --- | --- | --- | --- | --- |
| NULL | -655.4849 | NA | NA | NA |
| Larval_food | -654.8291 | 1.311576 | 1 | 2.52E-01 |
| Adult_food | -644.9383 | 19.78161 | 1 | 8.68E-06 |
| Larval_food: Adult_food | -643.7258 | 2.424919 | 1 | 1.19E-01 |
| ST2c: Females Late age) | | | | |
|  | **Loglik** | **Chisq** | **Df** | **Pr(>\|Chi\|)** |
| NULL | -645.3408 | NA | NA | NA |
| Larval_food | -644.8664 | 0.948705 | 1 | 0.3300493 |
| Adult_food | -643.5792 | 2.574501 | 1 | 0.1085984 |
| Larval_food: Adult_food | -643.4604 | 0.23748 | 1 | 0.6260325 |

**ST2d:** Males (Early age)

|  | Loglik | Chisq | Df | Pr(>\|Chi\|) |
| --- | --- | --- | --- | --- |
| NULL | -655.4849 | NA | NA | NA |
| Larval_food | -654.4296 | 2.11060654 | 1 | 0.146281 |
| Adult_food | -654.4184 | 0.02226572 | 1 | 0.881382 |
| Larval_food: Adult_food | -653.6421 | 1.55261894 | 1 | 0.212749 |

**ST2e:** Males (Mid age)

|  | Loglik | Chisq | Df | | Pr(>\|Chi\|) |
| --- | --- | --- | --- | --- | --- |
| NULL | -645.3408 | NA | | NA | NA |
| Larval_food | -645.3335 | 0.01451101 | | 1 | 0.904117 |
| Adult_food | -645.2042 | 0.25861141 | | 1 | 0.611076 |
| Larval_food: Adult_food | -643.3397 | 3.72903861 | | 1 | 0.053474 |

**ST2f:** Males (Late age)

|  | Loglik | Chisq | Df | Pr(>\|Chi\|) |
| --- | --- | --- | --- | --- |
| NULL | -655.4849 | NA | NA | NA |
| Larval_food | -655.4406 | 0.08846125 | 1 | 7.66E-01 |
| Adult_food | -645.9358 | 19.00971602 | 1 | 1.30E-05 |
| Larval_food: Adult_food | -643.3295 | 5.21262023 | 1 | 2.24E-02 |

**ST2g:** Pairwise comparisons

|  | Contrast | estimate | SE | df | z. ratio | p.value |
| --- | --- | --- | --- | --- | --- | --- |
| 1 | **(CC) - (PC)** | -0.20586 | 0.22813 | Inf | -0.9023787 | 8.04E-01 |
| 2 | **(CC) - (CP)** | -1.1217 | 0.237402 | Inf | -4.7249102 | 1.37E-05 |
| 3 | **(CC) - (PP)** | -0.58853 | 0.229589 | Inf | -2.5634173 | 5.07E-02 |
| 4 | **(PC) - (CP)** | -0.91585 | 0.232025 | Inf | -3.9471867 | 4.59E-04 |
| 5 | **(PC) - (PP)** | -0.38267 | 0.225951 | Inf | -1.6936145 | 3.27E-01 |
| 6 | **(CP) - (PP)** | 0.533172 | 0.22729 | Inf | 2.3457787 | 8.79E-02 |

**Supplementary table ST3:**

**Results of age and sex stratified analysis for Osmotic stress resistance (OSR)**

(model: coxme(Surv(Time, Status)~Larval_food*Adult_food +(1|Vial), data = data))

**ST3a:** Females (Early age)

|  | Df | Chisq | Pr(>Chisq) |
| --- | --- | --- | --- |
| Larval_food | 1 | 4.489941 | 0.034095 |
| Adult_food | 1 | 5.809079 | 0.015944 |
| Larval_food: Adult_food | 1 | 0.405456 | 0.524285 |

**ST3b:** Females (Mid-age)

|  | Df | Chisq | Pr(>Chisq) |
| --- | --- | --- | --- |
| Larval_food | 1 | 22.08676 | 2.61E-06 |
| Adult_food | 1 | 32.7592 | 1.04E-08 |
| Larval_food: Adult_food | 1 | 0.426317 | 5.14E-01 |

**ST3c:** Females (Late age)

|  | Df | Chisq | Pr(>Chisq) |
| --- | --- | --- | --- |
| Larval_food | 1 | 1.586083 | 0.207887 |
| Adult_food | 1 | 7.963253 | 0.004774 |
| Larval_food: Adult_food | 1 | 0.074182 | 0.785342 |

**ST3d:** Males (Early age)

|  | Df | Chisq | Pr(>Chisq) |
| --- | --- | --- | --- |
| Larval_food | 1 | 0.005501 | 0.940875 |
| Adult_food | 1 | 1.26881 | 0.25999 |
| Larval_food: Adult_food | 1 | 1.791244 | 0.180775 |

**ST3e:** Males (Mid age)

|  | Df | Chisq | Pr(>Chisq) |
| --- | --- | --- | --- |
| Larval_food | 1 | 8.61877 | 3.33E-03 |
| Adult_food | 1 | 100.3234 | 1.29E-23 |
| Larval_food: Adult_food | 1 | 2.899453 | 8.86E-02 |

**ST3f:** Males (Late age)

|  | Df | Chisq | Pr(>Chisq) |
| --- | --- | --- | --- |
| Larval_food | 1 | 0.003908 | 9.50E-01 |
| Adult_food | 1 | 21.37328 | 3.78E-06 |
| Larval_food: Adult_food | 1 | 8.631839 | 3.30E-03 |

**Supplementary table ST4:**

**Results of age and sex stratified analysis for Heat resistance (HR).**

(model: coxph(Surv(Time, Status)~Larval_food*Adult_food, data = data))

**ST4a:** Females (Early age)

|  | Loglik | Chisq | Df | Pr(>\|Chi\|) |
| --- | --- | --- | --- | --- |
| NULL | -496.4055 | NA | NA | NA |
| Larval_food | -485.225 | 22.360929 | 1 | 2.26E-06 |
| Adult_food | -482.9624 | 4.525201 | 1 | 3.34E-02 |
| Larval_food: Adult_food | -473.6833 | 18.558144 | 1 | 1.65E-05 |

**ST4b:** Pairwise comparison

|  | Contrast | estimate | SE | df | z.ratio | p.value |
| --- | --- | --- | --- | --- | --- | --- |
| 1 | **(CC) - (PC)** | -0.1487065 | 0.254175 | Inf | -0.58506 | 9.37E-01 |
| 2 | **(CC) - (CP)** | 1.3288662 | 0.291068 | Inf | 4.565489 | 2.95E-05 |
| 3 | **(CC) - (PP)** | -0.4777237 | 0.255878 | Inf | -1.867 | 2.42E-01 |
| 4 | **(PC) - (CP)** | 1.4775727 | 0.303606 | Inf | 4.86674 | 6.76E-06 |
| 5 | **(PC) - (PP)** | -0.3290172 | 0.253391 | Inf | -1.29846 | 5.64E-01 |
| 6 | **(CP) - (PP)** | -1.8065899 | 0.305865 | Inf | -5.9065 | 2.09E-08 |

**ST4c:** Females (Mid-age)

|  | Loglik | Chisq | Df | Pr(>\|Chi\|) |
| --- | --- | --- | --- | --- |
| NULL | -496.4055 | NA | NA | NA |
| Larval_food | -494.8072 | 3.196459 | 1 | 7.38E-02 |
| Adult_food | -472.3941 | 44.826292 | 1 | 2.15E-11 |
| Larval_food: Adult_food | -472.1568 | 0.474592 | 1 | 4.91E-01 |

**ST4d:** Females (Late age)

|  | Loglik | Chisq | Df | Pr(>\|Chi\|) |
| --- | --- | --- | --- | --- |
| NULL | -496.4055 | NA | NA | NA |
| Larval_food | -495.8641 | 1.082819 | 1 | 2.98E-01 |
| Adult_food | -475.8176 | 40.092948 | 1 | 2.42E-10 |
| Larval_food: Adult_food | -474.7492 | 2.136781 | 1 | 1.44E-01 |

**ST4e:** Males (Early age)

|  | Loglik | Chisq | Df | Pr(>\|Chi\|) |
| --- | --- | --- | --- | --- |
| NULL | -496.4055 | NA | NA | NA |
| Larval_food | -494.6324 | 3.546085 | 1 | 5.97E-02 |
| Adult_food | -467.4756 | 54.313769 | 1 | 1.71E-13 |
| Larval_food: Adult_food | -454.2878 | 26.375574 | 1 | 2.81E-07 |

**ST4f:** Pairwise comparison

|  | Contrast | Estimate | SE | df | z. ratio | p.value |
| --- | --- | --- | --- | --- | --- | --- |
| 1 | **(CC) - (PC)** | 0.8959083 | 0.262024 | Inf | 3.419187 | 3.51E-03 |
| 2 | **(CC) - (CP)** | 2.955302 | 0.348753 | Inf | 8.47392 | 3.87E-14 |
| 3 | **(CC) - (PP)** | 1.8514953 | 0.29495 | Inf | 6.27732 | 2.07E-09 |
| 4 | **(PC) - (CP)** | 2.0593937 | 0.327305 | Inf | 6.291977 | 1.88E-09 |
| 5 | **(PC) - (PP)** | 0.9555869 | 0.271411 | Inf | 3.520812 | 2.43E-03 |
| 6 | **(CP) - (PP)** | -1.1038067 | 0.283724 | Inf | -3.89042 | 5.79E-04 |

**ST4g:** Males (Mid age)

|  | Loglik | Chisq | Df | Pr(>\|Chi\|) |
| --- | --- | --- | --- | --- |
| NULL | -496.4055 | NA | NA | NA |
| Larval_food | -496.4015 | 0.0078767 | 1 | 0.92928 |
| Adult_food | -492.5036 | 7.7959204 | 1 | 0.005236 |
| Larval_food: Adult_food | -492.2997 | 0.4076957 | 1 | 0.523141 |

**ST4h:** Males (Late age)

|  | Loglik | Chisq | Df | Pr(>\|Chi\|) |
| --- | --- | --- | --- | --- |
| NULL | -491.5534 | NA | NA | NA |
| Larval_food | -486.9123 | 9.282346 | 1 | 0.002314 |
| Adult_food | -481.1332 | 11.558148 | 1 | 0.000675 |
| Larval_food: Adult_food | -480.4349 | 1.396657 | 1 | 0.237284 |

**Supplementary table ST5:**

**Results of age- and sex stratified analysis for Cold resistance (CR).**

model <- glm (status ~ Larval_food * Adult_food, family = binomial(link = "logit"), data=data)

**ST5a:** Females (Early age)

|  | Df | Deviance | Resid. Df | Resid. Dev | Pr(>Chi) |
| --- | --- | --- | --- | --- | --- |
| NULL | NA | NA | 129 | 96.98287 | NA |
| Larval_food | 1 | 5.033536 | 128 | 91.94934 | 0.024861 |
| Adult_food | 1 | 0.296808 | 127 | 91.65253 | 0.585891 |
| Larval_food: Adult_food | 1 | 0.115028 | 126 | 91.5375 | 0.734491 |

**ST5b:** Females (mid-age)

|  | Df | Deviance | Resid. Df | Resid. Dev | Pr(>Chi) |
| --- | --- | --- | --- | --- | --- |
| NULL | NA | NA | 133 | 165.0447 | NA |
| Larval_food | 1 | 0.3164995 | 132 | 164.7282 | 5.74E-01 |
| Adult_food | 1 | 45.6377648 | 131 | 119.0904 | 1.42E-11 |
| Larval_food: Adult_food | 1 | 1.9685609 | 130 | 117.1218 | 1.61E-01 |

**ST5c:** Females (late age)

|  | Df | Deviance | Resid. Df | Resid. Dev | Pr(>Chi) |
| --- | --- | --- | --- | --- | --- |
| NULL | NA | NA | 132 | 181.6536 | NA |
| Larval_food | 1 | 0.090293 | 131 | 181.5633 | 7.64E-01 |
| Adult_food | 1 | 8.503308 | 130 | 173.06 | 3.55E-03 |
| Larval_food: Adult_food | 1 | 15.29476 | 129 | 157.7652 | 9.20E-05 |

**ST5d:** Males (Early age)

|  | LR Chisq | Df | Pr(>Chisq) |
| --- | --- | --- | --- |
| Larval_food | 7.446732 | 1 | 6.36E-03 |
| Adult_food | 21.3724 | 1 | 3.78E-06 |
| Larval_food: Adult_food | 29.97032 | 1 | 4.39E-08 |

**ST5e:**

contrast	estimate	SE	df	z. ratio	p.value

(CC) - (PC)	-1.635	0.643	Inf	-2.545	0.0533

(CC) - (CP)	-18.187	1153.051	Inf	-0.016	1.0000

(CC) - (PP)	0.128	0.506	Inf	0.253	0.9943

(PC) - (CP)	-16.551	1153.051	Inf	-0.014	1.0000

(PC) - (PP)	1.764	0.641	Inf	2.753	0.0301

(CP) - (PP)	18.315	1153.051	Inf	0.016	1.0000

**ST5f:** Males (Mid age)

|  | LR Chisq | Df | Pr(>Chisq) |
| --- | --- | --- | --- |
| Larval_food | 22.10619 | 1 | 2.58E-06 |
| Adult_food | 21.67833 | 1 | 3.22E-06 |
| Larval_food:Adult_food | 24.65862 | 1 | 6.84E-07 |

| contrast estimate SE df z.ratio p.value |
| --- |
| (C-rich C-rich) - (P-rich C-rich) 20.14 2996.980 Inf 0.007 1.0000 |
| (C-rich C-rich) - (C-rich P-rich) 20.14 3040.733 Inf 0.007 1.0000 |
| (C-rich C-rich) - (P-rich P-rich) 1.33 0.601 Inf 2.207 0.1211 |
| (P-rich C-rich) - (C-rich P-rich) 0.00 4269.420 Inf 0.000 1.0000 |
| (P-rich C-rich) - (P-rich P-rich) -18.81 2996.980 Inf -0.006 1.0000 |
| (C-rich P-rich) - (P-rich P-rich) -18.81 3040.733 Inf -0.006 1.0000 |

**ST5g:** Males (Late age)

|  | LR Chisq | Df | Pr(>Chisq) |
| --- | --- | --- | --- |
| Larval_food | 22.10619 | 1 | 2.58E-06 |
| Adult_food | 21.67833 | 1 | 3.22E-06 |
| Larval_food:Adult_food | 24.65862 | 1 | 6.84E-07 |

**ST5h:**

|  | Contrast | Estimate | SE | df | z. ratio | p.value |
| --- | --- | --- | --- | --- | --- | --- |
| 1 | **(CC) - (PC)** | -2.3906 | 0.621354 | Inf | -3.8474 | 0.00068949 |
| 2 | **(CC) - (CP)** | -0.80256 | 0.518597 | Inf | -1.54757 | 0.409010156 |
| 3 | **(CC) - (PP)** | 1.526746 | 0.717859 | Inf | 2.126806 | 0.144564235 |
| 4 | **(PC) - (CP)** | 1.588034 | 0.599821 | Inf | 2.647514 | 0.04045556 |
| 5 | **(PC) - (PP)** | 3.917342 | 0.778565 | Inf | 5.03149 | 0.000002904 |
| 6 | **(CP) - (PP)** | 2.329308 | 0.699304 | Inf | 3.330897 | 0.00480085 |

**Supplementary table ST6:**

**Results of age- and sex stratified analysis for Desiccation resistance (DR).**

Desiccation resistance (DR): (model: coxme(Surv(Time,Status)~Larval_food*Adult_food +(1|Vial), data = data))

**ST6a:** Females (Early age)

|  | Df | Chisq | Pr(>Chisq) |
| --- | --- | --- | --- |
| Larval_food | 1 | 149.6578 | 2.06E-34 |
| Adult_food | 1 | 103.8924 | 2.14E-24 |
| Larval_food: Adult_food | 1 | 5.132149 | 2.35E-02 |

**ST6b:**

| Contrast | Estimate | SE | df | z. ratio | p.value |
| --- | --- | --- | --- | --- | --- |
| (CC) - (PC) | 2.13 | 0.193 | Inf | 11.030 | <.0001 |
| (CC) - (CP) | -1.15 | 0.174 | Inf | 6.586 | <.0001 |
| (CC) - (PP) | 0.44 | 0 .167 | Inf | 2.642 | 0.0410 |
| (PC) - (CP) | -3.28 | 0.228 | Inf | 14.379 | <.0001 |
| (PC) - (PP) | -1.69 | 0.182 | Inf | 9.271 | <.0001 |
| (CP) - (PP) | 1.59 | 0.183 | Inf | 8.663 | <.0001 |

**ST6c:** Females (Mid age)

|  | Df | Chisq | Pr(>Chisq) |
| --- | --- | --- | --- |
| Larval_food | 1 | 33.6089 | 6.74E-09 |
| Adult_food | 1 | 86.84708 | 1.17E-20 |
| Larval_food: Adult_food | 1 | 129.9657 | 4.17E-30 |

**ST6d:**

|  | Contrast | estimate | SE | df | z. ratio | p.value |
| --- | --- | --- | --- | --- | --- | --- |
| 1 | **(CC) - (PC)** | 0.417518 | 0.1660313 | Inf | 2.514697 | 0.0576615 |
| 2 | **(CC) - (CP)** | 3.476831 | 0.2365264 | Inf | 14.69955 | <.0001 |
| 3 | **(CC) - (PP)** | 0.724611 | 0.1713319 | Inf | 4.229282 | 0.0001377 |
| 4 | **(PC) - (CP)** | 3.059313 | 0.2263824 | Inf | 13.51392 | <.0001 |
| 5 | **(PC) - (PP)** | 0.307093 | 0.1635259 | Inf | 1.877945 | 0.2375206 |
| 6 | **(CP) - (PP)** | -2.75222 | 0.2183086 | Inf | -12.607 | <.0001 |

**ST6e:** Females (Late age)

|  | Df | Chisq | Pr(>Chisq) |
| --- | --- | --- | --- |
| Larval_food | 1 | 20.08042 | 7.43E-06 |
| Adult_food | 1 | 1.370024 | 2.42E-01 |
| Larval_food: Adult_food | 1 | 2.075375 | 1.50E-01 |

**ST6f:** Males (Early age)

|  | Df | Chisq | Pr(>Chisq) |
| --- | --- | --- | --- |
| Larval_food | 1 | 5.630698 | 1.76E-02 |
| Adult_food | 1 | 3.334518 | 6.78E-02 |
| Larval_food: Adult_food | 1 | 53.18751 | 3.03E-13 |

**ST6g:** Pairwise comparison

|  | contrast | Estimate | SE | df | z. ratio | p.value |
| --- | --- | --- | --- | --- | --- | --- |
| 1 | **(CC) - (PC)** | -1.27236 | 0.1797698 | Inf | -7.07771 | 8.822E-12 |
| 2 | **(CC) - (CP)** | -0.6083 | 0.1654288 | Inf | -3.67713 | 0.00134781 |
| 3 | **(CC) - (PP)** | -0.06668 | 0.1637688 | Inf | -0.40716 | 0.977208141 |
| 4 | **(PC) - (CP)** | 0.664054 | 0.1735247 | Inf | 3.826854 | 0.000748756 |
| 5 | **(PC - (PP)** | 1.205677 | 0.1809803 | Inf | 6.661927 | 1.62158E-10 |
| 6 | **(CP) - (PP)** | 0.541624 | 0.1623573 | Inf | 3.336 | 0.00471611 |

**ST6h:** Male (Mid age)

|  | Df | Chisq | Pr(>Chisq) |
| --- | --- | --- | --- |
| Larval_food | 1 | 103.5222 | 2.57E-24 |
| Adult_food | 1 | 23.82629 | 1.05E-06 |
| Larval_food: Adult_food | 1 | 9.780922 | 1.76E-03 |

**ST6i:** Pairwise comparison

|  | contrast | estimate | SE | df | z. ratio | p.value |
| --- | --- | --- | --- | --- | --- | --- |
| 1 | **(CC) - (PC)** | -1.81364 | 0.192893 | Inf | -9.40231 | 3.71E-14 |
| 2 | **(CC) - (CP)** | 0.269467 | 0.159968 | Inf | 1.684502 | 0.331842 |
| 3 | **(CC) - (PP)** | -0.79772 | 0.166747 | Inf | -4.78403 | 1.02E-05 |
| 4 | **(PC) - (CP)** | 2.083104 | 0.200122 | Inf | 10.4092 | 4.31E-14 |
| 5 | **(PC) - (PP)** | 1.015917 | 0.181672 | Inf | 5.592039 | 1.34E-07 |
| 6 | **(CP) - (PP)** | -1.06719 | 0.170346 | Inf | -6.26483 | 2.24E-09 |

**ST6j:** Males (Late age)

|  | Df | Chisq | Pr(>Chisq) |
| --- | --- | --- | --- |
| Larval_food | 1 | 5.068739 | 0.024361106 |
| Adult_food | 1 | 12.7626 | 0.000353619 |
| Larval_food: Adult_food | 1 | 1.712846 | 0.190616588 |

**Supplementary table ST7:**

**Results of age- and sex stratified analysis for Dry body weight.**

Dry Body weight: model <- glmer(Body weight ~ Larval_food * Adult_food + (1 | Vial), family = “gaussian”,(link = "identity"), data=data)

**ST7a:** Dry Body weight (Female Early age)

|  | LR Chisq | Df | Pr(>Chisq) |
| --- | --- | --- | --- |
| Larval_food | 10.1598506 | 1 | 1.44E-03 |
| Adult_food | 0.6419494 | 1 | 4.23E-01 |
| Larval_food: Adult_food | 30.5655954 | 1 | 3.23E-08 |

**ST7b:**

| contrast | estimate | SE | df | t. ratio | | p.value |
| --- | --- | --- | --- | --- | --- | --- |
| (CC) - (PC) | -0.0183 | 0.00574 | 36 | -3.187 | 0.0150 | |
| (CC) - (CP) | -0.0046 | 0.00574 | 36 | -0.801 | 0.8534 | |
| (CC) - (PP) | -0.0678 | 0.00574 | 36 | -11.807 | <.0001 | |
| (PC) - (CP) | 0.0137 | 0.00574 | 36 | 2.386 | 0.0980 | |
| (CP) - (PP) | -0.0632 | 0.00574 | 36 | -11.006 | <.0001 | |

**ST7c:** (Female mid age)

|  | LR Chisq | Df | Pr(>Chisq) |
| --- | --- | --- | --- |
| Larval_food | 14.9480217 | 1 | 1.11E-04 |
| Adult_food | 42.3530852 | 1 | 7.62E-11 |
| Larval_food: Adult_food | 0.2231139 | 1 | 6.37E-01 |

**ST7d:** Dry Body weight (Female late age)

|  | LR Chisq | Df | Pr(>Chisq) |
| --- | --- | --- | --- |
| Larval_food | 0.08697093 | 1 | 0.768064 |
| Adult_food | 14.86606397 | 1 | 0.000115 |
| Larval_food: Adult_food | 3.01606596 | 1 | 0.082443 |

**ST7e:** Dry body weight (Males Early age)

|  | LR Chisq | Df | Pr(>Chisq) |
| --- | --- | --- | --- |
| Larval_food | 5.655511 | 1 | 1.74E-02 |
| Adult_food | 33.82778 | 1 | 6.02E-09 |
| Larval_food: Adult_food | 56.73476 | 1 | 4.99E-14 |

**ST7f:**

| contrast | estimate | SE | df | t. ratio | p .value |
| --- | --- | --- | --- | --- | --- |
| CC-PC | 0.00696 | 0.00292 | 36 | 2.378 | 0.0996 |
| CC-CP | 0.01701 | 0.00292 | 36 | 5.816 | <.0001 |
| CC-PP | -0.00719 | 0.00292 | 36 | -2.458 | 0.0842 |
| PC-CP | 0.01006 | 0.00292 | 36 | 3.438 | 0.0078 |
| PC-PP | -0.01414 | 0.00292 | 36 | -4.836 | 0.0001 |
| CP-PP | -0.02420 | 0.00292 | 36 | -8.274 | <.0001 |

**ST7g:** Dry body weight (Males, mid-age)

|  | LR Chisq | Df | Pr(>Chisq) |
| --- | --- | --- | --- |
| Larval_food | 0.491114 | 1 | 4.83E-01 |
| Adult_food | 36.43183 | 1 | 1.58E-09 |
| Larval_food: Adult_food | 0.012776 | 1 | 9.10E-01 |

**ST7h:** Dry body weight (Males, Late age)

|  | LR Chisq | Df | Pr(>Chisq) |
| --- | --- | --- | --- |
| Larval_food | 0.10017 | 1 | 0.751626 |
| Adult_food | 0.10017 | 1 | 0.751626 |
| Larval_food: Adult_food | 1.645167 | 1 | 0.199618 |

**Supplementary table ST8:**

**Results of age-stratified analysis for Triglycerides (TAG).**

Triglycerides (TAG): glmer(TAG ~ Larval_food * Adult_food + (1 | Vial), family = “gaussian”,(link = "identity"), data=data)

**ST8a**: (Females, early age)

|  | Chisq | Df | Pr(>Chisq) |
| --- | --- | --- | --- |
| Larval_food | 0.002839 | 1 | 0.957509377 |
| Adult_food | 14.14156 | 1 | 0.000169556 |
| Larval_food: Adult_food | 0.306757 | 1 | 0.579676829 |

**ST8b:** TAG (Females, mid-age)

|  | Chisq | Df | Pr(>Chisq) |
| --- | --- | --- | --- |
| (Intercept) | 462.0229 | 1 | 1.74E-102 |
| Larval_food | 3.827644 | 1 | 5.04E-02 |
| Adult_food | 10.57702 | 1 | 1.15E-03 |
| Larval_food: Adult_food | 5.603336 | 1 | 1.79E-02 |

**ST8c**

| Contrast | null. Value | Estimate | std. error | Df | statistic | p.value |
| --- | --- | --- | --- | --- | --- | --- |
| CC - PC | 0 | -0.130355277 | 0.066629 | 42 | -1.95644 | 0.220792 |
| CC - CP | 0 | 0.21669279 | 0.066629 | 42 | 3.252233 | 0.011675 |
| CC - PP | 0 | 0.309386973 | 0.066629 | 42 | 4.643433 | 0.000191 |
| PC - CP | 0 | 0.347048067 | 0.066629 | 42 | 5.208669 | 3.12E-05 |
| PC - PP | 0 | 0.43974225 | 0.066629 | 42 | 6.59987 | 3.21E-07 |
| CP - PP | 0 | 0.092694183 | 0.066629 | 42 | 1.3912 | 0.511778 |

**ST8d:** TAG (Females, late age)

|  | Chisq | Df | Pr(>Chisq) |
| --- | --- | --- | --- |
| Larval_food: Adult_food | 3.57452 | 1 | 5.87E-02 |
| Adult_food | 30.70975 | 1 | 3.00E-08 |
| Larval_food: Adult_food | 2.16319 | 1 | 1.41E-01 |

**Supplementary table ST9:**

**Results of age-stratified analysis for Protein.**

Protein: glmer(protein ~ Larval_food * Adult_food + (1 | Vial), family = “gaussian”,(link = "identity"), data=data)

**ST9a:** (Female, early age)

|  | Chisq | Df | Pr(>Chisq) |
| --- | --- | --- | --- |
| (Intercept) | 215.1489 | 1 | 1.03E-48 |
| Larval_food | 26.57852 | 1 | 2.53E-07 |
| Adult_food | 50.35113 | 1 | 1.29E-12 |
| Larval_food: Adult_food | 14.16023 | 1 | 1.68E-04 |

**ST9b:** Pairwise comparison

| Contrast | null. value | estimate | std. error | Df | statistic | adj.p.value |
| --- | --- | --- | --- | --- | --- | --- |
| CC - PC | 0 | -0.088402808 | 0.017167 | 41.64027 | -5.14954 | 3.85E-05 |
| CC - CP | 0 | -0.119415321 | 0.016829 | 41.0371 | -7.09585 | 7.11E-08 |
| CC - PP | 0 | -0.117408142 | 0.016829 | 41.0371 | -6.97658 | 1.05E-07 |
| PC - CP | 0 | -0.031012513 | 0.017167 | 41.64027 | -1.80651 | 0.284775 |
| P C -PP | 0 | -0.029005334 | 0.017167 | 41.64027 | -1.68959 | 0.34193 |
| C P -PP | 0 | 0.002007178 | 0.016829 | 41.0371 | 0.11927 | 0.999383 |

**ST9c:** (Female mid-age)

|  | Chisq | Df | Pr(>Chisq) |
| --- | --- | --- | --- |
| (Intercept) | 303.2187 | 1 | 6.55E-68 |
| Larval_food | 8.364315 | 1 | 3.83E-03 |
| Adult_food | 81.53438 | 1 | 1.72E-19 |
| Larval_food: Adult_food | 10.79369 | 1 | 1.02E-03 |

**ST9d:** Pairwise comparison

| Contrast | null. value | estimate | std.error | Df | statistic | adj.p.value |
| --- | --- | --- | --- | --- | --- | --- |
| CC - PC | 0 | -0.03574 | 0.012356 | 42 | -2.89211 | 0.029615 |
| CC - CP | 0 | -0.11157 | 0.012356 | 42 | -9.02964 | 1.30E-10 |
| CC - PP | 0 | -0.0899 | 0.012356 | 42 | -7.27553 | 3.49E-08 |
| PC - CP | 0 | -0.07584 | 0.012356 | 42 | -6.13753 | 1.48E-06 |
| PC - PP | 0 | -0.05416 | 0.012356 | 42 | -4.38342 | 0.000431 |
| CP - PP | 0 | 0.021674 | 0.012356 | 42 | 1.75411 | 0.309502 |

**ST9e:** Protein (Female late age)

|  | Chisq | Df | Pr(>Chisq) |
| --- | --- | --- | --- |
| (Intercept) | 312.2591 | 1 | 7.03E-70 |
| Larval_food | 0.172302 | 1 | 6.78E-01 |
| Adult_food | 52.73681 | 1 | 3.81E-13 |
| Larval_food: Adult_food | 4.130057 | 1 | 4.21E-02 |

**ST9f:** Pairwise comparison

| Contrast | null. value | estimate | std. error | df | statistic | p.value |
| --- | --- | --- | --- | --- | --- | --- |
| CC – PC | 0 | -0.00469 | 0.011293 | 42 | -0.41509 | 0.975589 |
| CC – CP | 0 | -0.08201 | 0.011293 | 42 | -7.26201 | 3.64E-08 |
| CC – PP | 0 | -0.05424 | 0.011293 | 42 | -4.80306 | 0.000115 |
| PC – CP | 0 | -0.07732 | 0.011293 | 42 | -6.84692 | 1.42E-07 |
| PC – PP | 0 | -0.04956 | 0.011293 | 42 | -4.38797 | 0.000425 |
| CP – PP | 0 | 0.02777 | 0.011293 | 42 | 2.458949 | 0.081541 |

**Supplementary table ST10:**

**Results of age- and sex stratified analysis for Dry weight loss.**

Dry weight loss: glmer(Weight loss ~ Larval_food * Adult_food + (1 | Vial), family = “gaussian”,(link = "identity"), data=data)

**ST10a:** (Female, early age )

|  | Chisq | Df | Pr(>Chisq) |
| --- | --- | --- | --- |
| Larval_food | 14.57382 | 1 | 0.000135 |
| Adult_food | 14.8259 | 1 | 0.000118 |
| Larval_food: Adult_food | 0.688408 | 1 | 0.406706 |

**ST10b:** (Female, mid-age)

|  | Chisq | Df | Pr(>Chisq) |
| --- | --- | --- | --- |
| Larval_food | 1.5754 | 1 | 2.09E-01 |
| Adult_food | 53.60602 | 1 | 2.45E-13 |
| Larval_food: Adult_food | 0.109807 | 1 | 7.40E-01 |

**ST10c:** (Female, late age)

|  | Sum Sq | Df | F value | Pr(>F) |
| --- | --- | --- | --- | --- |
| Larval_food | 0.002157 | 1 | 0.823944 | 0.370066 |
| Adult_food | 0.000674 | 1 | 0.257344 | 0.615045 |
| Larval_food: Adult_food | 0.005886 | 1 | 2.248483 | 0.142465 |
| Residuals | 0.094241 | 36 | NA | NA |

**ST10d:** (Male, early age)

|  | Chisq | Df | Pr(>Chisq) |
| --- | --- | --- | --- |
| (Intercept) | 5227.792 | 1 | 0.00E+00 |
| Larval_food | 19.24382 | 1 | 1.15E-05 |
| Adult_food | 0.632179 | 1 | 4.27E-01 |
| Larval_food: Adult_food | 11.83561 | 1 | 5.81E-04 |

**ST10e:** (Male, mid-age)

|  | Chisq | Df | Pr(>Chisq) |
| --- | --- | --- | --- |
| Larval_food | 0.298181 | 1 | 0.585025 |
| Adult_food | 13.63575 | 1 | 0.000222 |
| Larval_food: Adult_food | 1.752733 | 1 | 0.185533 |

**ST10f:** (Male, late age)

|  | Chisq | Df | Pr(>Chisq) |
| --- | --- | --- | --- |
| (Intercept) | 698.0909 | 1 |  |
| Larval_food | 3.344624 | 1 | 6.74E-02 |
| Adult_food | 2.2443 | 1 | 1.34E-01 |
| Larval_food: Adult_food | 4.277453 | 1 | 3.86E-02 |

**ST10g:** Pairwise comparison

| Contrast | null. value | Estimate | std. error | Df | statistic | adj.p.value |
| --- | --- | --- | --- | --- | --- | --- |
| CC - PC | 0 | 0.040261 | 0.022015 | 27 | 1.828831 | 0.282252 |
| CC - CP | 0 | 0.03298 | 0.022015 | 27 | 1.498099 | 0.452493 |
| CC – PP | 0 | 0.008851 | 0.022015 | 27 | 0.402053 | 0.977563 |
| PC – CP | 0 | -0.00728 | 0.022015 | 27 | -0.33073 | 0.98724 |
| PC – PP | 0 | -0.03141 | 0.022015 | 27 | -1.42678 | 0.49422 |
| CP – PP | 0 | -0.02413 | 0.022015 | 27 | -1.09605 | 0.694914 |

**Supplementary table ST11:**

**Results of age- and sex stratified analysis for Water in control condition.**

**Water Control:** glmer(waterC ~ Larval_food * Adult_food + (1 | Vial), family = “gaussian”,(link = "identity"), data=data)

**ST11a:** (Females, Early age)

|  | Sum Sq | Df | F value | Pr(>F) |
| --- | --- | --- | --- | --- |
| Larval_food | 1.28E-03 | 1 | 4.853789 | 0.03407 |
| Adult_food | 1.74E-03 | 1 | 6.587295 | 0.014575 |
| Larval_food: Adult_food | 3.86E-05 | 1 | 0.146241 | 0.704402 |
| Residuals | 9.51E-03 | 36 | NA | NA |

**ST11b:** (Females, mid-age)

|  | Chisq | Df | Pr(>Chisq) |
| --- | --- | --- | --- |
| Larval_food | 0.586281 | 1 | 4.44E-01 |
| Adult_food | 16.87239 | 1 | 4.00E-05 |
| Larval_food: Adult_food | 0.108237 | 1 | 7.42E-01 |

**ST11c:** (Females, late-age)

|  | Sum Sq | Df | Pr(>F) |
| --- | --- | --- | --- |
| Larval_food | 1.42E-02 | 1 | 4.64E-08 |
| Adult_food | 2.26E-03 | 1 | 9.25E-03 |
| Larval_food: Adult_food | 1.20E-03 | 1 | 5.24E-02 |

**ST11d:** (Males, early age)

|  | Chisq | Df | Pr(>Chisq) |
| --- | --- | --- | --- |
| (Intercept) | 1.94E+05 | 1 | 0.00E+00 |
| Larval_food | 7.06E-01 | 1 | 4.01E-01 |
| Adult_food | 2.42E+01 | 1 | 8.91E-07 |
| Larval_food: Adult_food | 1.93E+01 | 1 | 1.13E-05 |

**ST11e:** Pairwise comparison

| contrast | null. value | estimate | std.error | Df | statistic | adj.p.value |
| --- | --- | --- | --- | --- | --- | --- |
| CC - PC | 0 | -0.00251 | 0.002995 | 25.22113 | -0.83887 | 0.835469 |
| CC - CP | 0 | -0.0147 | 0.002995 | 25.22113 | -4.90676 | 0.000256 |
| CC – PP | 0 | 0.00158 | 0.002995 | 25.22113 | 0.527702 | 0.951562 |
| PC – CP | 0 | -0.01218 | 0.003061 | 24.46759 | -3.9797 | 0.002836 |
| PC – PP | 0 | 0.004093 | 0.003061 | 24.46759 | 1.33695 | 0.549188 |
| CP – PP | 0 | 0.016276 | 0.003061 | 24.46759 | 5.316646 | 9.81E-05 |

**ST11f:** (Male, mid-age)

|  | LR Chisq | Df | Pr(>Chisq) |
| --- | --- | --- | --- |
| Larval_food | 0.017752 | 1 | 0.894005 |
| Adult_food | 2.134204 | 1 | 0.144045 |
| Larval_food: Adult_food | 1.094912 | 1 | 0.295386 |

**ST11g:** (Male, late age)

|  | Sum Sq | Df | F value | Pr(>F) |
| --- | --- | --- | --- | --- |
| Larval_food | 5.24E-03 | 1 | 11.09574 | 0.00205 |
| Adult_food | 1.73E-06 | 1 | 0.003654 | 0.952144 |
| Larval_food: Adult_food | 8.05E-04 | 1 | 1.702555 | 0.200469 |
| Residuals | 1.65E-02 | 35 | NA | NA |

**Supplementary table ST12:**

**Results of age- and sex stratified analysis for Water under osmotic stress condition.**

Water OSR: glmer(WaterOSR ~ Larval_food * Adult_food + (1 | Vial), family = “gaussian”,(link = "identity"), data=data)

**ST12a:** Early females

|  | Sum Sq | Df | F value | Pr(>F) |
| --- | --- | --- | --- | --- |
| (Intercept) | 8.408926 | 1 | 43895.76 | 3.67E-57 |
| Larval_food | 0.001235 | 1 | 6.44834 | 1.56E-02 |
| Adult_food | 0.00407 | 1 | 21.24539 | 4.93E-05 |
| Larval_food: Adult_food | 0.000805 | 1 | 4.200634 | 4.77E-02 |
| Residuals | 0.006896 | 36 | NA | NA |

**ST12b:** Pairwise comparison

| contrast | null. value | estimate | std.error | Df | statistic | adj.p.value |
| --- | --- | --- | --- | --- | --- | --- |
| CC - PC | 0 | 0.015718 | 0.00619 | 36 | 2.539358 | 0.070609 |
| CC - CP | 0 | -0.02853 | 0.00619 | 36 | -4.60927 | 0.000277 |
| CC – PP | 0 | -0.03075 | 0.00619 | 36 | -4.96841 | 9.43E-05 |
| PC – CP | 0 | -0.04425 | 0.00619 | 36 | -7.14863 | 1.24E-07 |
| PC – PP | 0 | -0.04647 | 0.00619 | 36 | -7.50777 | 4.23E-08 |
| CP – PP | 0 | -0.00222 | 0.00619 | 36 | -0.35914 | 0.98388 |

**ST12c:** (Females, mid-age)

|  | Sum Sq | Df | F value | Pr(>F) |
| --- | --- | --- | --- | --- |
| Larval_food | 2.33E-03 | 1 | 8.026908 | 0.007505 |
| Adult_food | 4.33E-04 | 1 | 1.494115 | 0.229522 |
| Larval_food: Adult_food | 6.53E-05 | 1 | 0.225074 | 0.638066 |
| Residuals | 1.04E-02 | 36 | NA | NA |

**ST12d:** (Females, late age)

|  | Chisq | Df | Pr(>Chisq) |
| --- | --- | --- | --- |
| (Intercept) | 3.32E+04 | 1 | 0.00E+00 |
| Larval_food | 3.68E+01 | 1 | 1.28E-09 |
| Adult_food | 3.59E-01 | 1 | 5.49E-01 |
| Larval_food: Adult_food | 4.99E+01 | 1 | 1.58E-12 |

**ST12e:** Pairwise comparison

| contrast | null. value | estimate | Std. error | Df | statistic | adj.p.value |
| --- | --- | --- | --- | --- | --- | --- |
| CC - PC | 0 | -0.04125 | 0.006795 | 27 | -6.07041 | 1.01E-05 |
| CC - CP | 0 | -0.00407 | 0.006795 | 27 | -0.59897 | 0.931486 |
| CC – PP | 0 | 0.022592 | 0.006795 | 27 | 3.324957 | 0.012826 |
| PC – CP | 0 | 0.037177 | 0.006795 | 27 | 5.471447 | 4.86E-05 |
| PC – PP | 0 | 0.063839 | 0.006795 | 27 | 9.395371 | 3.12E-09 |
| CP – PP | 0 | 0.026662 | 0.006795 | 27 | 3.923924 | 0.002867 |

**ST12f:** (Males, early age)

|  | Sum Sq | Df | F value | Pr(>F) |
| --- | --- | --- | --- | --- |
| Larval_food | 1.40E-03 | 1 | 5.761203 | 2.22E-02 |
| Adult_food | 7.94E-03 | 1 | 32.63592 | 2.25E-06 |
| Larval_food: Adult_food | 3.96E-06 | 1 | 0.016277 | 8.99E-01 |
| Residuals | 8.03E-03 | 33 | NA | NA |

**ST12g:** (Males, mid-age)

|  | Chisq | Df | Pr(>Chisq) |
| --- | --- | --- | --- |
| Larval_food | 3.349175 | 1 | 0.067239 |
| Adult_food | 6.538351 | 1 | 0.010557 |
| Larval_food: Adult_food | 1.250555 | 1 | 0.263447 |

**ST12h:** (Males, late age)

|  | Sum Sq | Df | F value | Pr(>F) |
| --- | --- | --- | --- | --- |
| Larval_food | 0.001223 | 1 | 1.704697 | 0.200192 |
| Adult_food | 0.006056 | 1 | 8.441442 | 0.006319 |
| Larval_food: Adult_food | 0.000229 | 1 | 0.319781 | 0.575347 |
| Residuals | 0.025109 | 35 | NA | NA |

**Supplementary table ST13:**

**Results of age- and sex stratified analysis for Water in desiccation condition.**

Water DR: glmer(waterDR ~ Larval_food * Adult_food + (1 | Vial), family = “gaussian”,(link = "identity"), data=data)

**ST13a:** (Females, early age)

|  | Sum Sq | Df | F value | Pr(>F) |
| --- | --- | --- | --- | --- |
| Larval_food | 0.000696 | 1 | 3.03138 | 9.07E-02 |
| Adult_food | 0.006393 | 1 | 27.84857 | 7.50E-06 |
| Larval_food: Adult_food | 0.000355 | 1 | 1.548682 | 2.22E-01 |
| Residuals | 0.007805 | 34 | NA | NA |

**ST13b:** (Females, mid-age)

|  | Sum Sq | Df | F value | Pr(>F) |
| --- | --- | --- | --- | --- |
| (Intercept) | 9.003241 | 1 | 28579.95 | 8.23E-54 |
| Larval_food | 0.005394 | 1 | 17.12368 | 2.01E-04 |
| Adult_food | 0.006258 | 1 | 19.86633 | 7.79E-05 |
| Larval_food: Adult_food | 0.003476 | 1 | 11.03365 | 2.06E-03 |
| Residuals | 0.011341 | 36 | NA | NA |

**ST13c:** Pairwise comparison

| contrast | null. value | estimate | std.error | df | statistic | adj.p.value |
| --- | --- | --- | --- | --- | --- | --- |
| CC - PC | 0 | -0.03285 | 0.007937 | 36 | -4.13808 | 0.001107 |
| CC - CP | 0 | -0.03538 | 0.007937 | 36 | -4.45717 | 0.000436 |
| CC – PP | 0 | -0.03094 | 0.007937 | 36 | -3.89766 | 0.002201 |
| PC – CP | 0 | -0.00253 | 0.007937 | 36 | -0.31909 | 0.98857 |
| PC – PP | 0 | 0.001908 | 0.007937 | 36 | 0.240418 | 0.995023 |
| CP – PP | 0 | 0.004441 | 0.007937 | 36 | 0.559507 | 0.943332 |

**ST13d:** (Females, late age)

|  | Sum Sq | Df | F value | Pr(>F) |
| --- | --- | --- | --- | --- |
| (Intercept) | 8.355889 | 1 | 22809.72 | 4.74E-52 |
| Larval_food | 0.004544 | 1 | 12.40422 | 1.18E-03 |
| Adult_food | 0.006398 | 1 | 17.46507 | 1.78E-04 |
| Larval_food: Adult_food | 0.00297 | 1 | 8.10686 | 7.24E-03 |
| Residuals | 0.013188 | 36 | NA | NA |

**ST13e:** Pairwise comparison

| contrast | null. value | estimate | std.error | df | statistic | adj.p.value |
| --- | --- | --- | --- | --- | --- | --- |
| CC - PC | 0 | -0.03015 | 0.00856 | 36 | -3.52196 | 0.006225 |
| CC - CP | 0 | -0.03577 | 0.00856 | 36 | -4.17912 | 0.000983 |
| CC – PP | 0 | -0.03145 | 0.00856 | 36 | -3.67446 | 0.004105 |
| PC – CP | 0 | -0.00563 | 0.00856 | 36 | -0.65716 | 0.912334 |
| PC – PP | 0 | -0.00131 | 0.00856 | 36 | -0.1525 | 0.998712 |
| CP – PP | 0 | 0.00432 | 0.00856 | 36 | 0.504664 | 0.95745 |

**ST13f: (**Males, early age)

|  | Chisq | Df | Pr(>Chisq) |
| --- | --- | --- | --- |
| Larval_food | 8.776695 | 1 | 0.003051 |
| Adult_food | 11.47568 | 1 | 7.05E-04 |
| Larval_food: Adult_food | 0.243077 | 1 | 6.22E-01 |

**ST13g: (**Males, mid-age)

|  | Sum Sq | Df | F value | Pr(>F) |
| --- | --- | --- | --- | --- |
| Larval_food | 0.000254 | 1 | 0.47251 | 0.49624 |
| Adult_food | 0.003252 | 1 | 6.039143 | 0.018941 |
| Larval_food: Adult_food | 0.000685 | 1 | 1.271198 | 0.267001 |
| Residuals | 0.019385 | 36 | NA | NA |

**ST13h: (**Males, late age)

|  | Sum Sq | Df | F value | Pr(>F) |
| --- | --- | --- | --- | --- |
| (Intercept) | 9.067182 | 1 | 1.76E+04 | 4.82E-50 |
| Larval_food | 1.13E-03 | 1 | 2.19E+00 | 1.47E-01 |
| Adult_food | 1.02E-04 | 1 | 1.99E-01 | 6.58E-01 |
| Larval_food: Adult_food | 4.52E-03 | 1 | 8.79E+00 | 5.34E-03 |
| Residuals | 0.018508 | 36 | NA | NA |

**ST13i:** Pairwise comparison

| contrast | null. value | estimate | std. error | Df | statistic | p.value |
| --- | --- | --- | --- | --- | --- | --- |
| CC - PC | 0 | 0.015019 | 0.01014 | 36 | 1.481172 | 0.459048 |
| CC - CP | 0 | -0.00452 | 0.01014 | 36 | -0.44611 | 0.969953 |
| CC – PP | 0 | -0.03202 | 0.01014 | 36 | -3.15805 | 0.016195 |
| PC – CP | 0 | -0.01954 | 0.01014 | 36 | -1.92728 | 0.234899 |
| PC – PP | 0 | -0.04704 | 0.01014 | 36 | -4.63922 | 0.000254 |
| CP – PP | 0 | -0.0275 | 0.01014 | 36 | -2.71194 | 0.047895 |

**ST14: Larval diet effects made a limited contribution to the coordinated dietary-effect architecture. (Significant associations (P < 0.05) are highlighted in red)**

**ST14a:**

| Early age Female | | | | | | | | |
| --- | --- | --- | --- | --- | --- | --- | --- | --- |
| Male | LF | SR | DR | HR | OR | CR | Longevity | Fertility |
|  | SR | _ | 0.012766259 | 0.008478009 | 0.10060813 | 0.072600361 | 0.58052821 | 0.001853462 |
|  | DR | 0.397782956 | _ | 0.007319347 | 0.098182667 | 0.070169464 | 0.58022264 | 0.001395377 |
|  | HR | 0.620544583 | 0.637685659 | _ | 0.090370715 | 0.062414582 | 0.57923748 | 0.00044109 |
|  | OR | 0.467572778 | 0.511846477 | 0.654893928 | _ | 0.163333482 | 0.59186584 | 0.071382421 |
|  | CR | 0.435586652 | 0.48745352 | 0.646543366 | 0.532810339 | _ | 0.58859547 | 0.044171683 |
|  | Longevity | 0.272309299 | 0.372778488 | 0.614415749 | 0.450372777 | 0.415044218 | _ | 0.5768197 |
|  | Fertility | 0.159249952 | 0.300725359 | 0.598925912 | 0.40304445 | 0.357317511 | 0.09787472 | _ |

**ST14b:**

|  | Mid age Female | | | | | | | |
| --- | --- | --- | --- | --- | --- | --- | --- | --- |
| Male | LF | SR | DR | HR | OR | CR | Longevity | Fertility |
|  | SR |  | 0.249272529 | 0.924461908 | 0.217300039 | 0.769884543 | 0.60781943 | 0.254704666 |
|  | DR | 0.85791941 |  | 0.924306357 | 0.024944445 | 0.765372294 | 0.58419969 | 0.073748564 |
|  | HR | 0.952380516 | 0.94949651 |  | 0.924273505 | 0.927776197 | 0.92532226 | 0.924312303 |
|  | OR | 0.8596627 | 0.275784079 | 0.949576083 |  | 0.764388637 | 0.5786658 | 0.031563727 |
|  | CR | 0.868412292 | 0.66353104 | 0.950017471 | 0.684875696 |  | 0.79123703 | 0.765549151 |
|  | Longevity | 0.858630976 | 0.1010124 | 0.949528679 | 0.353203422 | 0.672657282 |  | 0.58517902 |
|  | Fertility | 0.858229985 | 0.022432744 | 0.949510499 | 0.312485833 | 0.667590062 | 0.16148998 |  |

**ST14c:**

| Late Female | | | | | | | | |
| --- | --- | --- | --- | --- | --- | --- | --- | --- |
| Male | LF | SR | DR | HR | OR | CR | Longevity | Fertility |
|  | SR |  | 0.445889843 | 0.564829154 | 0.406400163 | 0.628946694 | 0.62995817 | 0.449004606 |
|  | DR | 0.70484081 |  | 0.527473299 | 0.301825441 | 0.606537635 | 0.60774243 | 0.367613661 |
|  | HR | 0.669458972 | 0.473174554 |  | 0.503506529 | 0.656848802 | 0.65764971 | 0.529420331 |
|  | OR | 0.727277015 | 0.643828348 | 0.575908527 |  | 0.592972491 | 0.59430532 | 0.30787569 |
|  | CR | 0.677301176 | 0.50311765 | 0.247710733 | 0.592073739 |  | 0.69165478 | 0.607667103 |
|  | Longevity | 0.672651116 | 0.485731772 | 0.187548018 | 0.582574908 | 0.281648723 |  | 0.608861614 |
|  | Fertility | 0.680167767 | 0.513342856 | 0.280493968 | 0.597809323 | 0.350139621 | 0.31066579 |  |

**ST15: Adult-diet effect architecture linking stress-resistance traits with longevity and fertility across sex and age.**

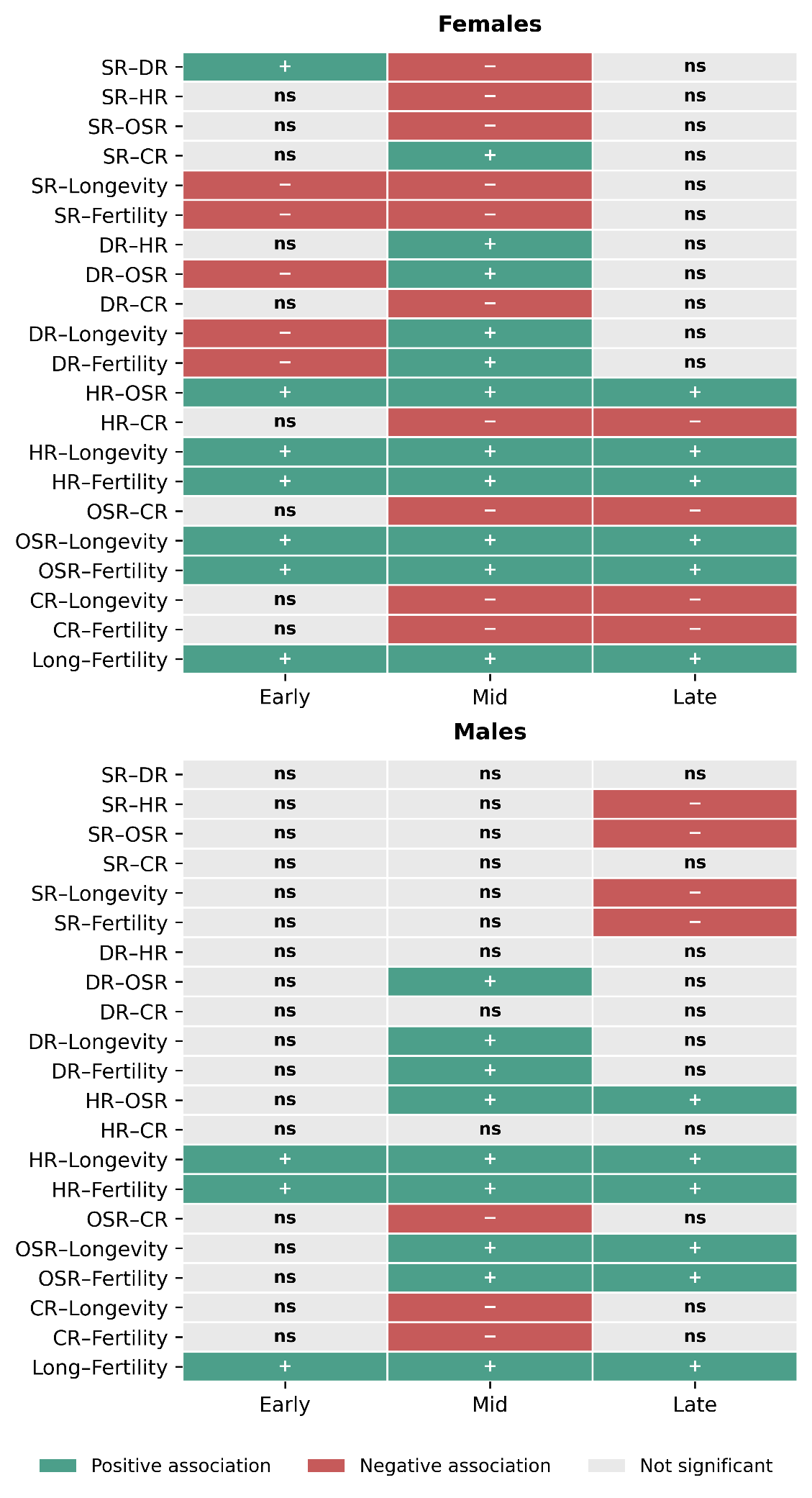

**ST16a:** **Z-score values for larval diet to test directional association at an early age**

|  | 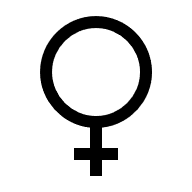 | | | | | | | |
| --- | --- | --- | --- | --- | --- | --- | --- | --- |
| 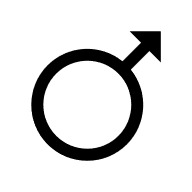 | **Larval diet** | **SR** | **DR** | **HR** | **OR** | **CR** | **Longevity** | **Fertility** |
|  | **SR** | - | 1.807881 | -0.57234 | -1.0246 | -0.09778 | -5.31812 | 0.370415 |
|  | **DR** | -2.22029 | - | -0.31658 | -0.56674 | -0.05408 | -2.94163 | 0.204889 |
|  | **HR** | -0.8571 | 0.386029 | - | 1.790209 | 0.170842 | 9.291952 | -0.6472 |
|  | **OR** | -1.51847 | 0.683909 | 1.771651 | - | 0.095431 | 5.190428 | -0.36152 |
|  | **CR** | -0.1364 | 0.061434 | 0.159144 | 0.089828 | - | 54.38926 | -3.78829 |
|  | **Longevity** | -0.90606 | 0.408081 | 1.057123 | 0.596688 | 6.642555 | - | -0.06965 |
|  | **Fertility** | 0.156518 | -0.07049 | -0.18261 | -0.10308 | -1.14747 | -0.17275 | - |

**ST16b:** **Z-score values for adult diet to test directional association at an early age**

|  | 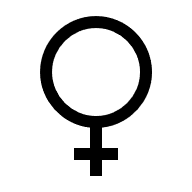 | | | | | | | |
| --- | --- | --- | --- | --- | --- | --- | --- | --- |
| 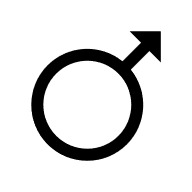 | **Adult diet** | **SR** | **DR** | **HR** | **OR** | **CR** | **Longevity** | **Fertility** |
|  | **SR** | - | 1.450643 | -0.55703 | -0.28208 | -0.26869 | -0.25328 | -0.27956 |
|  | **DR** | -0.2282 | - | -0.38399 | -0.19445 | -0.18522 | -0.1746 | -0.19271 |
|  | **HR** | -0.0053 | 0.023241 | - | 0.506395 | 0.482364 | 0.454693 | 0.501867 |
|  | **OR** | -0.18253 | 0.799837 | 34.41439 | - | 0.952545 | 0.897902 | 0.897902 |
|  | **CR** | -0.01964 | 0.086051 | 3.702506 | 0.107586 | - | 0.942634 | 1.040432 |
|  | **Longevity** | -0.02737 | 0.119932 | 5.160289 | 0.149946 | 1.393729 | - | 1.10375 |
|  | **Fertility** | -0.01529 | 0.067009 | 2.883161 | 0.083778 | 0.778705 | 0.558721 | - |

**ST16c:** **Z-score values for larval diet to test directional association at mid age**

|  | 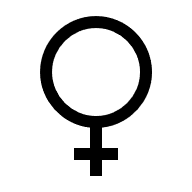 | | | | | | | | |
| --- | --- | --- | --- | --- | --- | --- | --- | --- | --- |
| 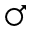 | **Larval diet** | | **SR** | **DR** | **HR** | **OR** | **CR** | **Longevity** | **Fertility** |
|  | **SR** | - | | -1.41834 | -10.6923 | -0.23016 | -0.40329 | -3.63384 | -0.22262 |
|  | **DR** | -0.04878 | | - | 7.538577 | 0.162271 | 0.28434 | -3.63384 | 0.156957 |
|  | **HR** | 1.176854 | | -24.1236 | - | 0.021525 | 0.037718 | 0.339857 | 0.02082 |
|  | **OR** | -0.05712 | | 1.170847 | -0.04854 | - | 1.752258 | 15.78865 | 0.967252 |
|  | **CR** | -0.00595 | | 0.121972 | -0.00506 | 0.104174 | - | 9.010458 | 0.552003 |
|  | **Longevity** | -0.11888 | | 2.436901 | -0.10102 | 2.081316 | 19.97914 | - | 0.061262 |
|  | **Fertility** | -0.01806 | | 0.370263 | -0.01535 | 0.316235 | 3.035629 | 0.15194 | - |

**ST16d:** **Z-score values for adult diet to test directional association at mid age**

|  | **Adult diet** | **SR** | **DR** | **HR** | **OR** | **CR** | **Longevity** | **Fertility** |
| --- | --- | --- | --- | --- | --- | --- | --- | --- |
|  | **SR** | - | -2.63618 | -0.4383 | -0.59414 | 0.089191 | -0.90026 | -0.20879 |
|  | **DR** | 0.186837 | - | 0.166265 | 0.225378 | -0.03383 | 0.341503 | 0.079204 |
|  | **HR** | 0.186837 | -0.15463 | - | 1.355537 | -0.20349 | 2.053968 | 0.47637 |
|  | **OR** | 0.039473 | 0.058513 | 0.211269 | - | -0.15012 | 1.515243 | 0.351425 |
|  | **CR** | -0.01959 | -0.02904 | -0.10485 | -0.49628 | - | -10.0936 | -2.34098 |
|  | **Longevity** | 0.2598 | 0.385116 | 1.390515 | 6.581722 | -13.262 | - | 0.231927 |
|  | **Fertility** | 0.030501 | 0.045213 | 0.163249 | 0.772707 | -1.55699 | 0.117402 | - |

**ST16e:** **Z-score values for larval diet to test directional association at late age**

|  | **LF** | **SR** | **DR** | **HR** | **OR** | **CR** | **Longevity** | **Fertility** |
| --- | --- | --- | --- | --- | --- | --- | --- | --- |
|  | **SR** |  | -1.04717 | 0.599712 | -0.48933 | 0.479976 | -2.15809 | -0.23897 |
|  | **DR** | 0.996754 |  | -0.5727 | 0.467288 | -0.45836 | 2.060884 | 0.22821 |
|  | **HR** | -0.14097 | -0.14143 |  | -0.81594 | 0.800345 | -3.59855 | -0.39848 |
|  | **OR** | 0.062344 | 0.062547 | -0.44224 |  | -0.98089 | 4.410306 | 0.488371 |
|  | **CR** | 0.191041 | 0.191663 | -1.35514 | 3.064279 |  | -4.49625 | -0.49789 |
|  | **Longevity** | -0.51285 | -0.51452 | 3.637889 | -8.22609 | -2.68451 |  | 0.110734 |
|  | **Fertility** | -0.14085 | -0.14131 | 0.999097 | -2.25918 | -0.73726 | 0.274636 |  |

**ST16f:** **Z-score values for adult diet to test directional association at late age**

|  | AF | SR | DR | HR | OR | CR | Longevity | Fertility |
| --- | --- | --- | --- | --- | --- | --- | --- | --- |
|  | SR |  | -2.17716 | -0.08251 | -0.33023 | 0.060615 | -0.208 | -0.06321 |
|  | DR | -6.19511 |  | 0.037899 | 0.15168 | -0.02784 | 0.09554 | 0.029034 |
|  | HR | -1.20081 | 0.193832 |  | 4.00225 | -0.73463 | 2.520923 | 0.766098 |
|  | OR | -0.60282 | 0.097306 | 0.502014 |  | -0.18355 | 0.629877 | 0.191417 |
|  | CR | 0.253615 | -0.04094 | -0.2112 | -0.42071 |  | -3.43156 | -1.04284 |
|  | Longevity | -1.87381 | 0.302466 | 1.560455 | 3.10839 | -7.3884 |  | 0.303896 |
|  | Fertility | -0.28825 | 0.046529 | 0.240049 | 0.478172 | -1.13658 | 0.153833 |  |

**ST17:** **RMST table**

| **Sex** | **Day** | **Regime** | **OSR** | **SR** | **HR** | **DR** |
| --- | --- | --- | --- | --- | --- | --- |
| F | Early | CC | 98.76 | 92.09 | 21.34 | 32.83 |
| F | Early | CP | 124.08 | 90.04 | 28.37 | 31.27 |
| F | Early | PC | 82.84 | 109.23 | 20.42 | 35.33 |
| F | Early | PP | 114.48 | 96.86 | 19.57 | 33.35 |
| F | Late | CC | 16.8 | 86.18 | 3.22 | 15.09 |
| F | Late | CP | 20.68 | 82.05 | 6.28 | 15.48 |
| F | Late | PC | 15.44 | 91.74 | 3.08 | 14.33 |
| F | Late | PP | 18.36 | 85.08 | 7.25 | 14.8 |
| F | Mid | CC | 30.68 | 82.82 | 6.08 | 19.3 |
| F | Mid | CP | 52.36 | 71.08 | 12.14 | 23.18 |
| F | Mid | PC | 23.64 | 99.77 | 6.58 | 19.8 |
| F | Mid | PP | 33.72 | 67.81 | 11.5 | 20.2 |
| M | Early | CC | 122.64 | 54.38 | 8.73 | 19.55 |
| M | Early | CP | 122.68 | 51.16 | 31.15 | 18.85 |
| M | Early | PC | 115.72 | 54.24 | 15.37 | 18.05 |
| M | Early | PP | 121.71 | 57 | 22.14 | 19.45 |
| M | Late | CC | 8.16 | 63.22 | 4.67 | 7.13 |
| M | Late | CP | 17.52 | 41.52 | 5.4 | 7.7 |
| M | Late | PC | 10.96 | 58.32 | 3.35 | 7.58 |
| M | Late | PP | 13.32 | 49.59 | 4.8 | 7.7 |
| M | Mid | CC | 21.2 | 52.76 | 6.63 | 10.98 |
| M | Mid | CP | 52.88 | 51.33 | 9.07 | 11.18 |
| M | Mid | PC | 16.04 | 49.56 | 7.43 | 9.1 |
| M | Mid | PP | 49.6 | 55.16 | 8.36 | 10.13 |

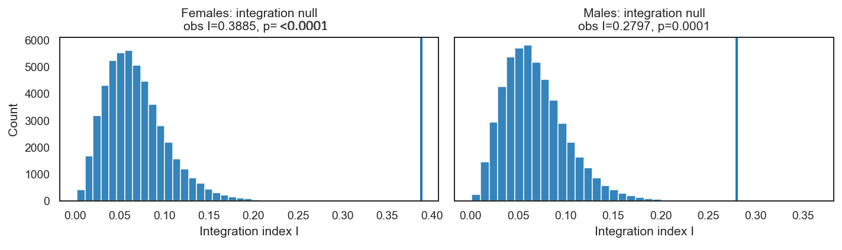

**Fig S1.** **Permutation tests of resistance-trait integration in females and males.** The integration index ($I$) was calculated separately for each sex from the eigenvalue distribution of the complete Pearson correlation matrix among five standardized resistance traits across 12 age × dietary-regime conditions. Null distributions were generated using 50,000 permutations in which condition-level values were shuffled independently within each trait, preserving each trait’s marginal distribution while disrupting correspondence among traits. Histograms show the resulting permutation distributions, and vertical lines indicate the observed integration indices. Observed integration exceeded all permuted values in both females ($I=0.43$, $P<0.0001$) and males ($I=0.3$, $P<0.0001$), indicating stronger condition-level integration among resistance traits than expected in the absence of systematic correspondence among traits. The numerically higher female index is descriptive because the analysis did not directly test the difference between sexes.

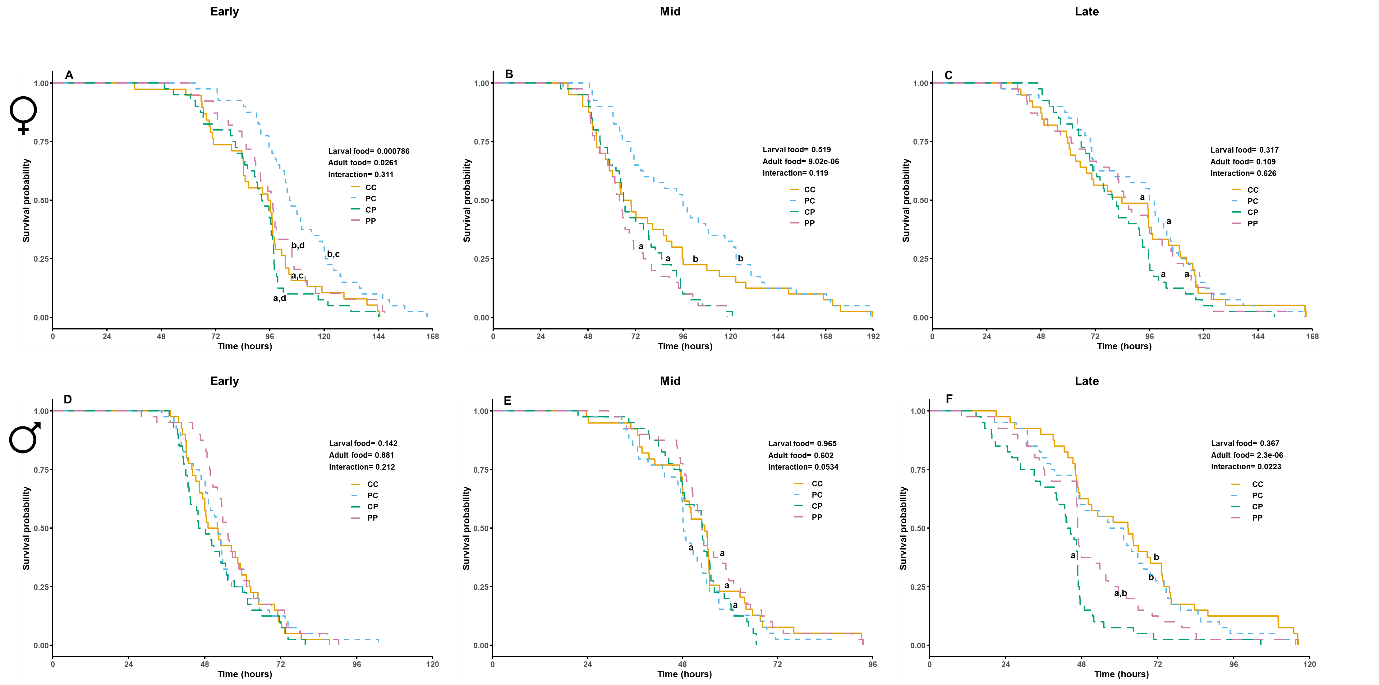

**Fig S2. Stage-specific nutrition modifies starvation resistance across age- and sex-dependent manner.** Survival curves illustrating dietary responses in early, mid and late age females (A, B, C) and males (D, E, F) for Starvation resistance. Starvation resistance in females, showing a significant larval and adult diet effect at early age (A), significant adult diet effect at mid age (B), and no significant effect at late age (C). In males there was no significant effect of diet on SR at both early (D) and mid (E) age but interaction between larval and adult diet at a late age. Diet regimes are denoted by two letters, where the first letter indicates larval diet and the second indicates adult diet: CC, carbohydrate-rich larval and adult diet; PC, protein-rich larval and carbohydrate-rich adult diet; CP, carbohydrate-rich larval and protein-rich adult diet; PP, protein-rich larval and adult diet. LD, larval diet; AD, adult diet. P values indicate model-based tests of larval diet, adult diet and their interaction within the corresponding sex–age stratum. Different letters indicate statistical differences among diet regimes.

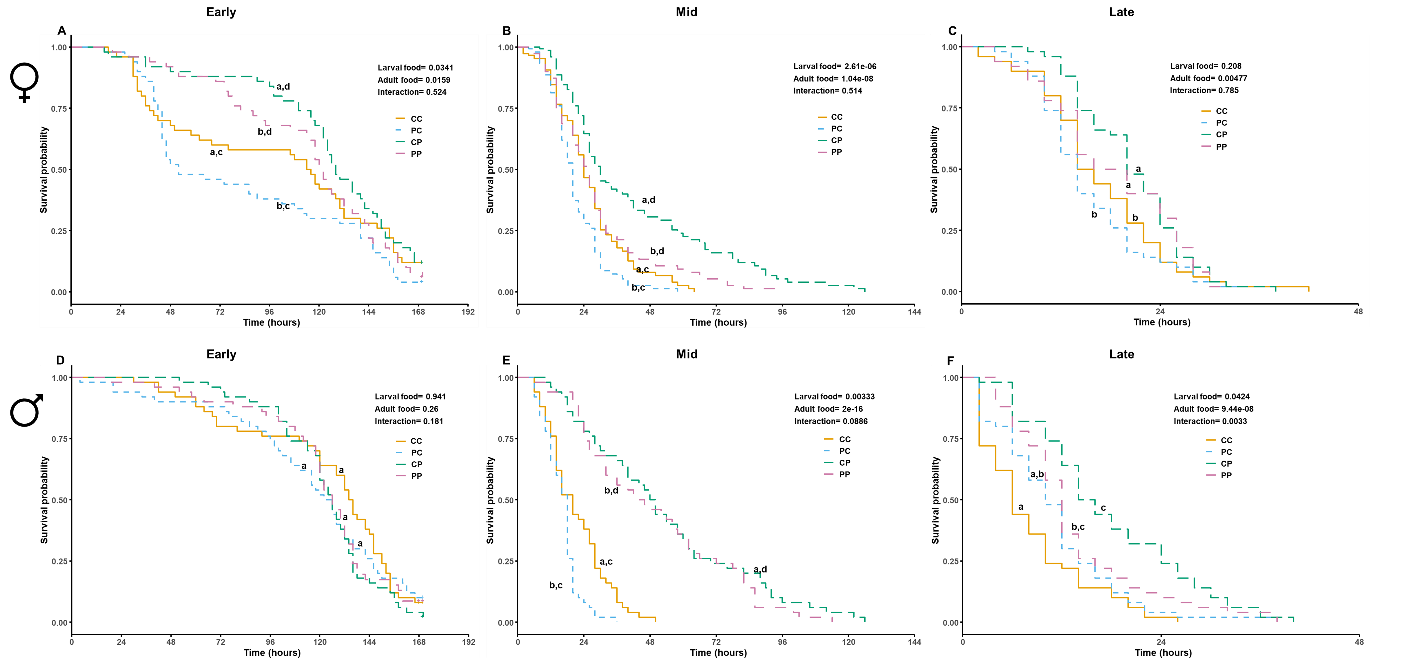

**Fig S3.** **Stage-specific nutrition modifies osmotic stress resistance across age- and sex-dependent manner.** Survival curves illustrating dietary responses in early, mid and late age females (A, B, C) and males (D, E, F) for Osmotic stress resistance (OSR). OSR in females, showing a significant larval and adult diet effect at early age (A) and mid age (B), and adult diet effect at late age (C). In males, there was no significant effect of diet on OSR at early age (D), both larval and adult diet effects at mid age (E), but an interaction between larval and adult diet at a late age. Diet regimes are denoted by two letters, where the first letter indicates larval diet and the second indicates adult diet: CC, carbohydrate-rich larval and adult diet; PC, protein-rich larval and carbohydrate-rich adult diet; CP, carbohydrate-rich larval and protein-rich adult diet; PP, protein-rich larval and adult diet. LD, larval diet; AD, adult diet. P values indicate model-based tests of larval diet, adult diet and their interaction within the corresponding sex–age stratum. Different letters indicate statistical differences among diet regimes.

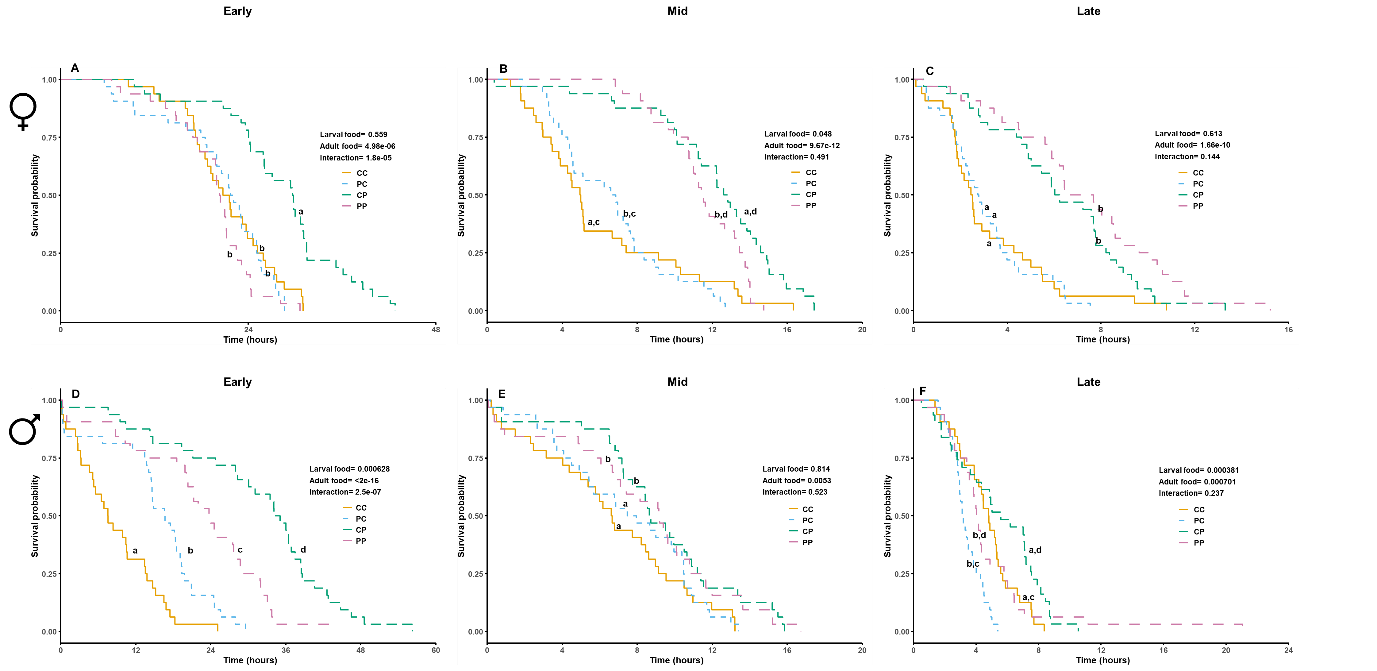

**Fig S4.** **Stage-specific nutrition modifies heat resistance across age- and sex-dependent manner.** Survival curves illustrating dietary responses in early, mid and late age females (A, B, C) and males (D, E, F) for Heat resistance (HR). Heat resistance in females, showing a significant interaction between larval and adult diet at early age (A), a significant larval and adult diet effect at mid age (B), and adult diet effect at late age (C). In males also there was interaction effect at early (D) age, adult diet effect at mid age (E) and both larval and adult diet effect at late age. Diet regimes are denoted by two letters, where the first letter indicates larval diet and the second indicates adult diet: CC, carbohydrate-rich larval and adult diet; PC, protein-rich larval and carbohydrate-rich adult diet; CP, carbohydrate-rich larval and protein-rich adult diet; PP, protein-rich larval and adult diet. LD, larval diet; AD, adult diet. P values indicate model-based tests of larval diet, adult diet and their interaction within the corresponding sex–age stratum. Different letters indicate statistical differences among diet regimes.

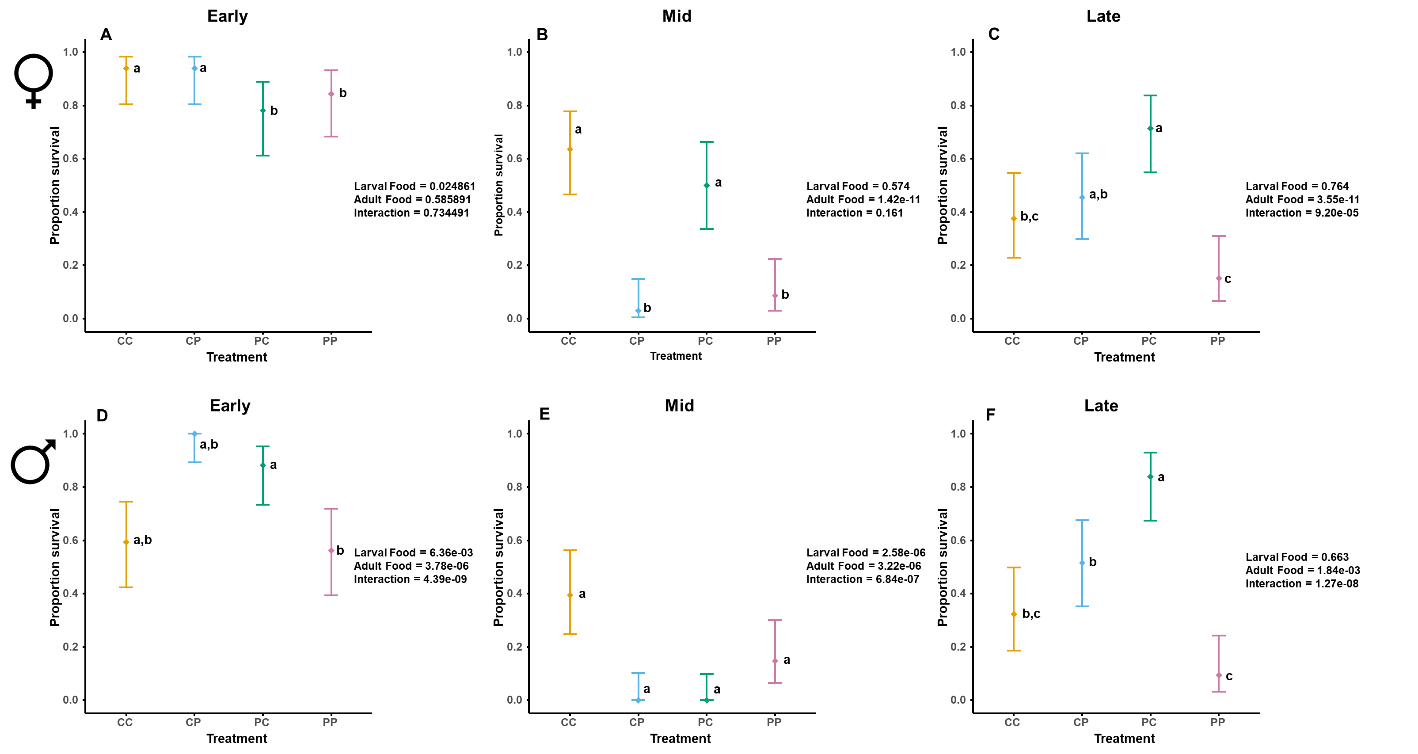

**Fig S5.** Dot plot indicating dietary responses in early, mid and late age females (A, B, C) and males (D, E, F) for Cold resistance (CR). CR in females, showing a significant larval diet effect at early age (A), a significant adult diet effect at mid age (B), and interaction between larval and adult diet effect at late age (C). In males, there was an interaction effect at all ages. Diet regimes are denoted by two letters, where the first letter indicates larval diet and the second indicates adult diet: CC, carbohydrate-rich larval and adult diet; PC, protein-rich larval and carbohydrate-rich adult diet; CP, carbohydrate-rich larval and protein-rich adult diet; PP, protein-rich larval and adult diet. LD, larval diet; AD, adult diet. P values indicate model-based tests of larval diet, adult diet and their interaction within the corresponding sex–age stratum. Different letters indicate statistical differences among diet regimes.

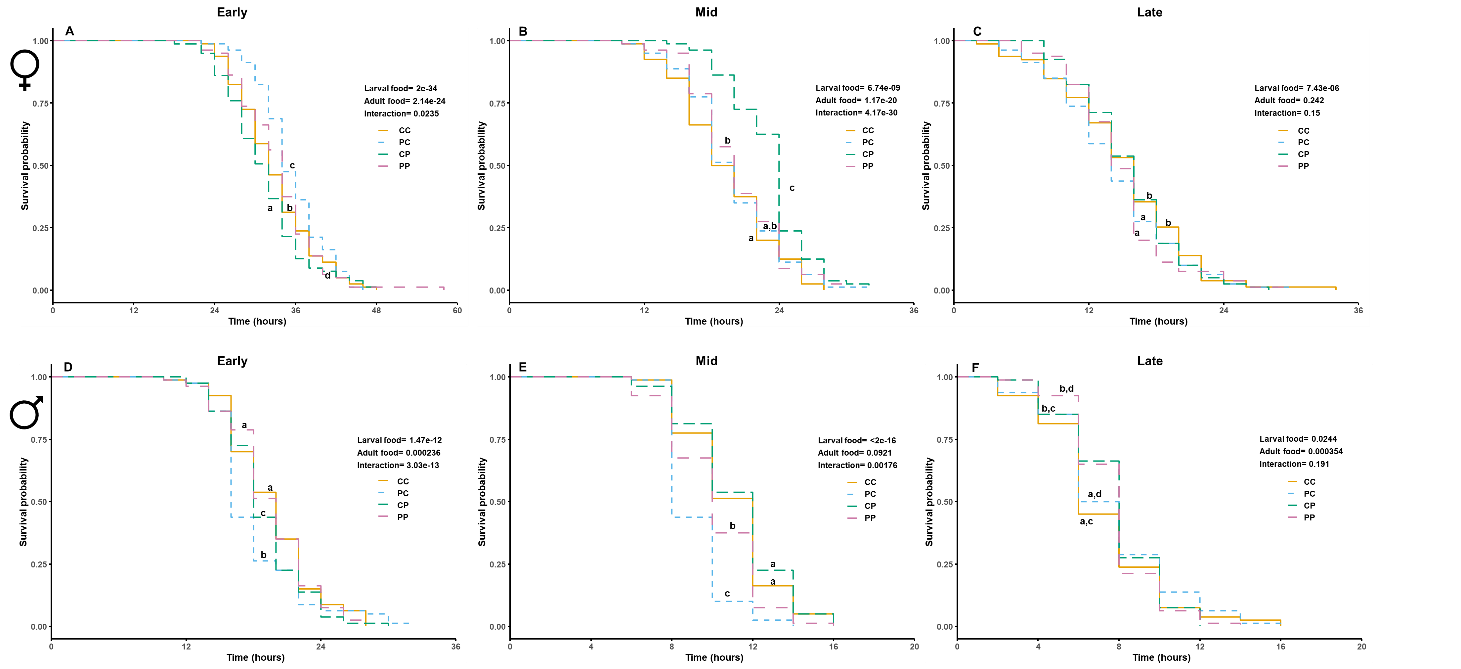

**Fig S6.** Survival curves illustrating dietary responses in early, mid and late age females (A, B, C) and males (D, E, F) for Desiccation resistance (DR). Desiccation resistance in females, showing a significant interaction between larval and adult diet at early (A), and mid age (B), and larval diet effect at late age (C). In males, there was also an interaction effect at early (D) and mid age (E), and both larval and adult diet effects at late age. Diet regimes are denoted by two letters, where the first letter indicates larval diet and the second indicates adult diet: CC, carbohydrate-rich larval and adult diet; PC, protein-rich larval and carbohydrate-rich adult diet; CP, carbohydrate-rich larval and protein-rich adult diet; PP, protein-rich larval and adult diet. LD, larval diet; AD, adult diet. P values indicate model-based tests of larval diet, adult diet and their interaction within the corresponding sex–age stratum. Different letters indicate statistical differences among diet regimes.

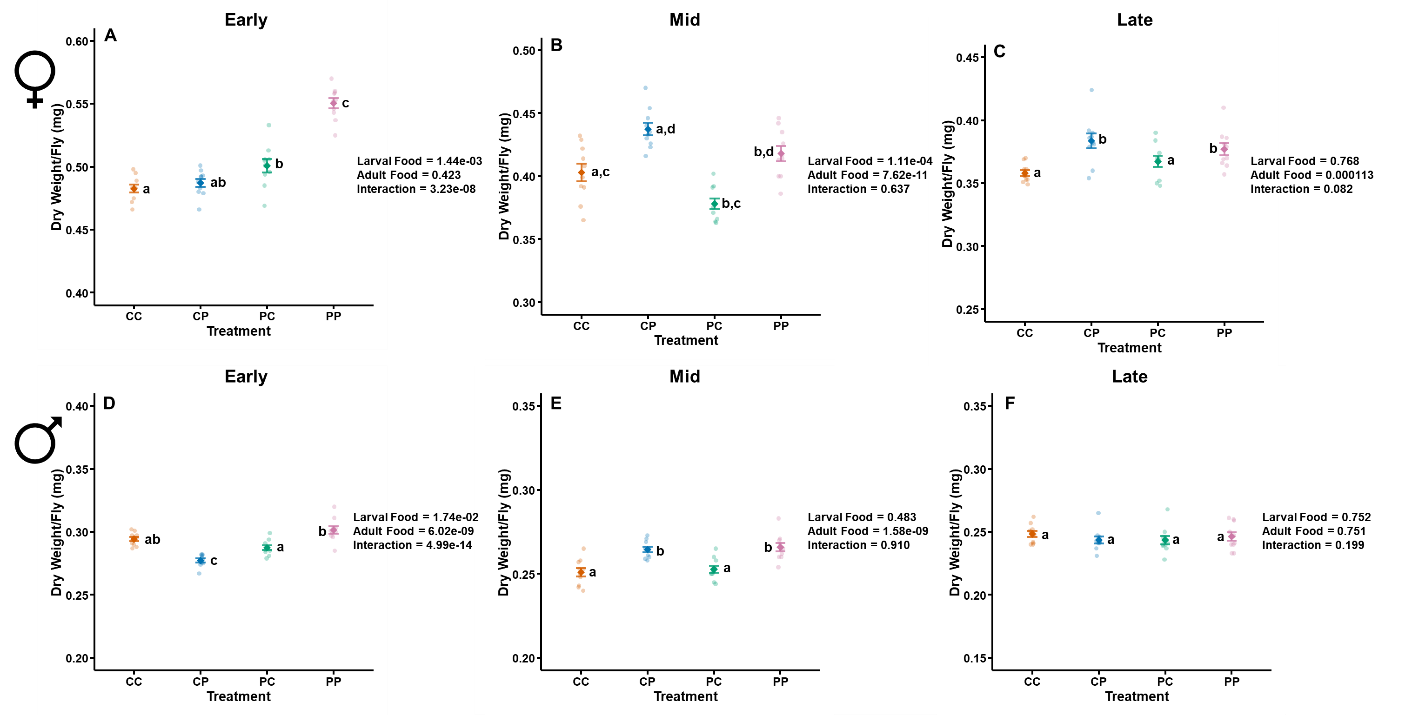

**Fig S7.** A–C, Whole-body dry weight across larval × adult diet regimes in early (A), mid (B) and late (C) adult females. D–F, Whole-body dry weight across the same diet regimes and ages in males. Coloured circles show biological replicates, and group means with error bars are shown in corresponding darker colours. Diet regimes are denoted by two letters, where the first letter indicates larval diet and the second indicates adult diet: CC, carbohydrate-rich larval and adult diet; CP, carbohydrate-rich larval and protein-rich adult diet; PC, protein-rich larval and carbohydrate-rich adult diet; PP, protein-rich larval and adult diet. P values indicate model-based tests of larval diet (LD), adult diet (AD) and their interaction (LD × AD). Different letters indicate significant statistical differences among diet regimes. Adult carbohydrate-rich diet was associated with elevated TAG, whereas adult protein-rich diet was associated with elevated protein concentration, indicating distinct adult-diet-dependent physiological states.

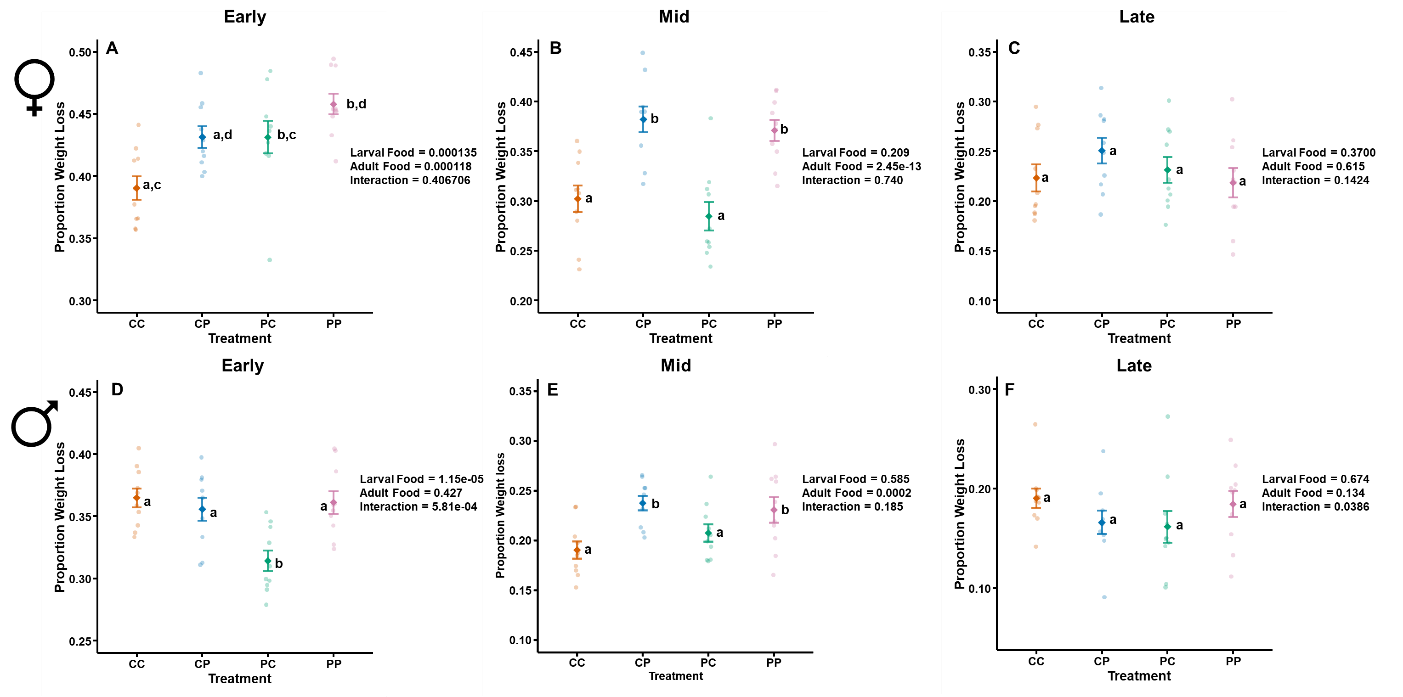

**Fig S8.** A–C, Dry weight loss under starvation across larval × adult diet regimes in early (A), mid (B) and late (C) adult females. D–F, Whole-body dry weight across the same diet regimes and ages in males. Coloured circles show biological replicates, and group means with error bars are shown in corresponding darker colours. Diet regimes are denoted by two letters, where the first letter indicates larval diet and the second indicates adult diet: CC, carbohydrate-rich larval and adult diet; CP, carbohydrate-rich larval and protein-rich adult diet; PC, protein-rich larval and carbohydrate-rich adult diet; PP, protein-rich larval and adult diet. P values indicate model-based tests of larval diet (LD), adult diet (AD) and their interaction (LD × AD). Different letters indicate significant statistical differences among diet regimes.

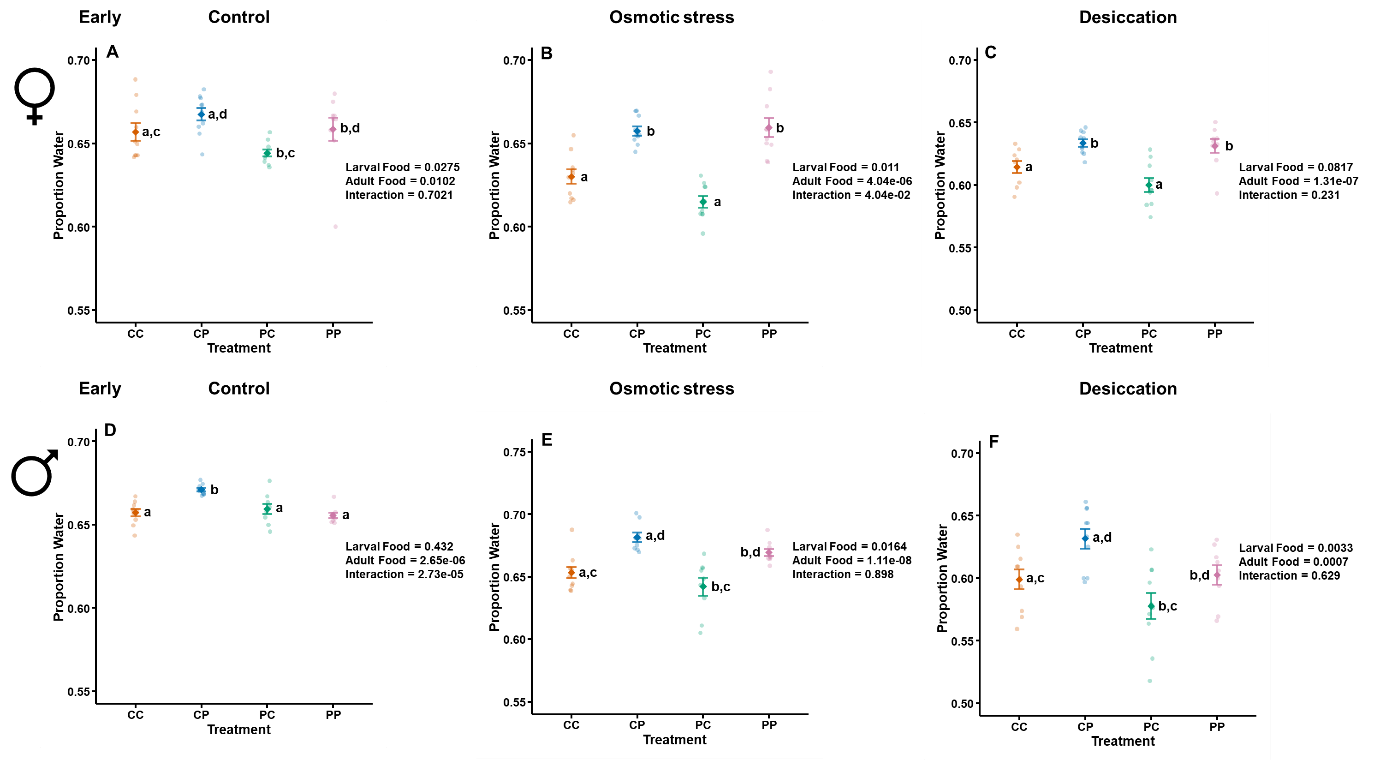

**Fig S9.** A–C, Proportion water across larval × adult diet regimes in control (A), osmotic stress (B) and Desiccation (C) adult females at early age. D–F, Proportion water across the same diet regimes and stress condition in males. Coloured circles show biological replicates, and group means with error bars are shown in corresponding darker colours. Diet regimes are denoted by two letters, where the first letter indicates larval diet and the second indicates adult diet: CC, carbohydrate-rich larval and adult diet; CP, carbohydrate-rich larval and protein-rich adult diet; PC, protein-rich larval and carbohydrate-rich adult diet; PP, protein-rich larval and adult diet. P values indicate model-based tests of larval diet (LD), adult diet (AD) and their interaction (LD × AD). Different letters indicate significant statistical differences among diet regimes.

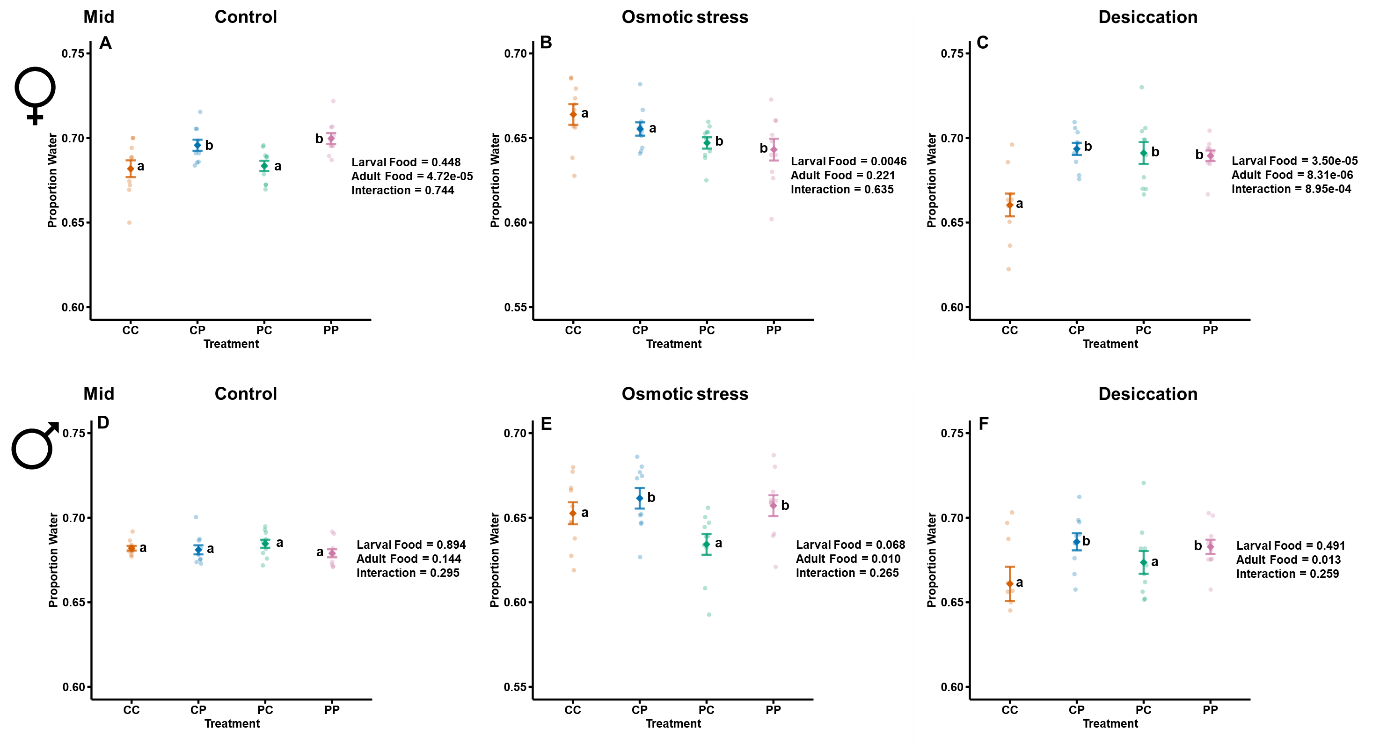

**Fig S10.** A–C, Proportion water across larval × adult diet regimes in control (A), osmotic stress (B) and Desiccation (C) adult females at mid age. D–F, Proportion water across the same diet regimes and stress condition in males. Coloured circles show biological replicates, and group means with error bars are shown in corresponding darker colours. Diet regimes are denoted by two letters, where the first letter indicates larval diet and the second indicates adult diet: CC, carbohydrate-rich larval and adult diet; CP, carbohydrate-rich larval and protein-rich adult diet; PC, protein-rich larval and carbohydrate-rich adult diet; PP, protein-rich larval and adult diet. P values indicate model-based tests of larval diet (LD), adult diet (AD) and their interaction (LD × AD). Different letters indicate significant statistical differences among diet regimes.

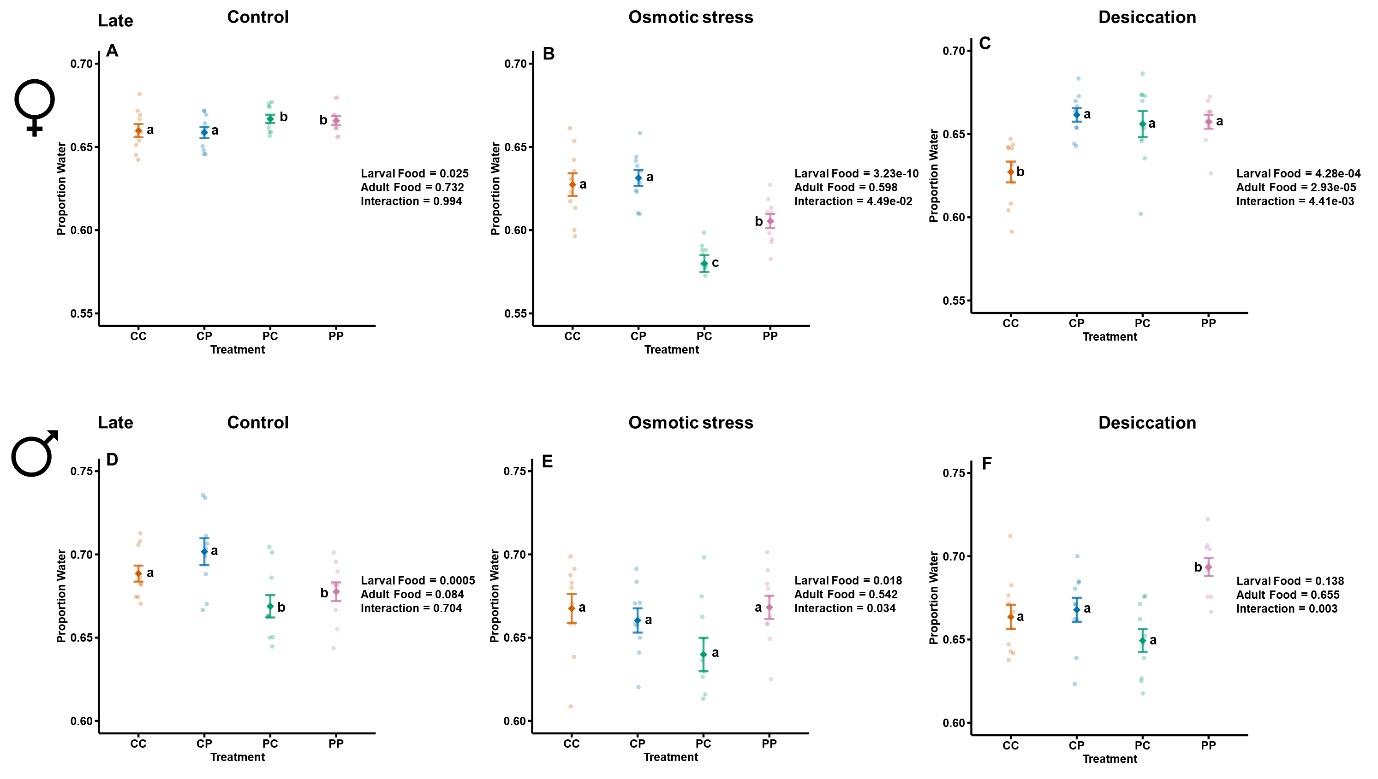

**Fig S11.** A–C, Proportion water across larval × adult diet regimes in control (A), osmotic stress (B) and Desiccation (C) adult females at late age. D–F, Proportion water across the same diet regimes and stress condition in males. Coloured circles show biological replicates, and group means with error bars are shown in corresponding darker colours. Diet regimes are denoted by two letters, where the first letter indicates larval diet and the second indicates adult diet: CC, carbohydrate-rich larval and adult diet; CP, carbohydrate-rich larval and protein-rich adult diet; PC, protein-rich larval and carbohydrate-rich adult diet; PP, protein-rich larval and adult diet. P values indicate model-based tests of larval diet (LD), adult diet (AD) and their interaction (LD × AD). Different letters indicate significant statistical differences among diet regimes.

| 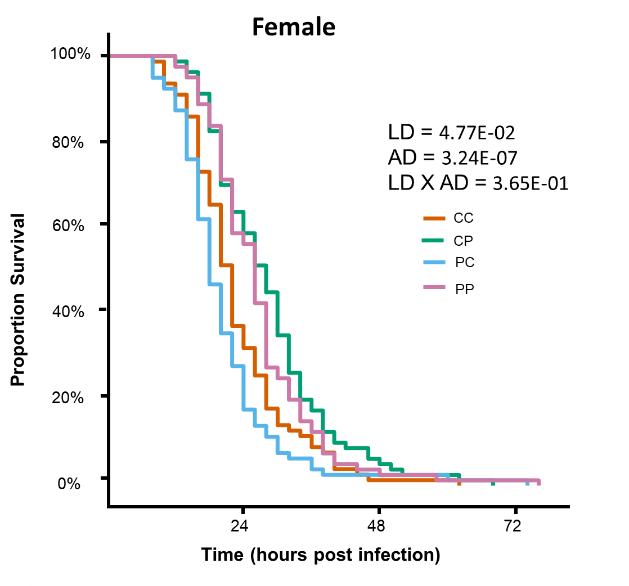 | 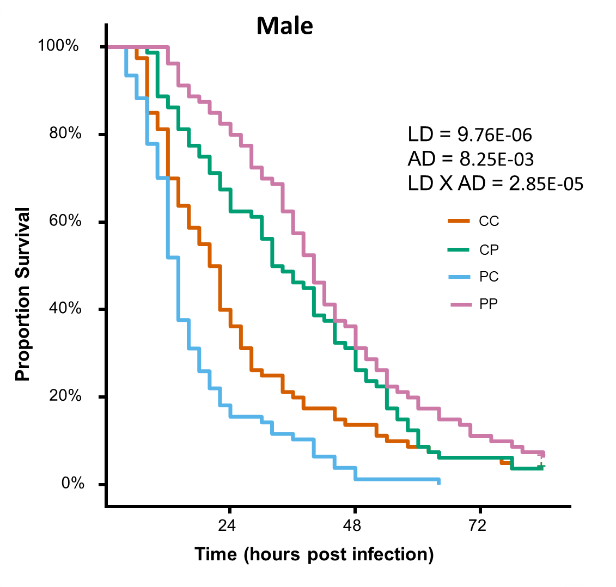 |
| --- | --- |
| **Fig S12.** **Stage-specific nutrition modifies osmotic stress resistance on Day 22 post egg collection in a sex-dependent manner.** Survival curves illustrating dietary responses in mid females (A) and males (B) for Osmotic stress resistance (OSR) under common food (P: C = 0.4). OSR in females, showing a significant larval and adult diet effect (A). In males, there was an interaction between larval and adult diet at mid-age (B). Diet regimes are denoted by two letters, where the first letter indicates larval diet and the second indicates adult diet: CC, carbohydrate-rich larval and adult diet; CP, carbohydrate-rich larval and protein-rich adult diet; PC, protein-rich larval and carbohydrate-rich adult diet; PP, protein-rich larval and adult diet. P values indicate model-based tests of larval diet (LD), adult diet (AD) and their interaction (LD × AD). Different letters indicate significant statistical differences among diet regimes. | |
